# SGEF coordinates epithelial morphogenesis by regulating junction stability, collective migration, and extracellular matrix remodeling

**DOI:** 10.64898/2026.07.17.739205

**Authors:** Madeline Lovejoy, Agustin Rabino, Lucia Gonzalez-Blotta, Vennela Gangasani, Sophia Durham, Gabriel Kreider, Rafael Garcia-Mata

## Abstract

Polarized epithelia are essential for organ function, and disruption of epithelial polarity is a hallmark of many diseases, including cancer. We previously showed that the RhoG-specific guanine nucleotide exchange factor SGEF interacts with the Scribble polarity complex to regulate epithelial junction assembly in 2D monolayers. However, its role in epithelial morphogenesis and lumen formation in 3D remains unknown. Here, we combined quantitative morphometric analysis with long-term live-cell imaging to investigate the role of SGEF during MDCK cyst development. SGEF KD disrupted normal lumenogenesis, producing enlarged cysts with multiple collapsed lumens accompanied by reduced E-cadherin, β-catenin, and ZO-1 expression. Loss of SGEF also altered the distribution of the actomyosin network. Re-expression of WT SGEF restored the normal phenotype, whereas restoration of E-cadherin and ZO-1 partially rescued lumen architecture, identifying the loss of junction integrity as a key driver of the morphogenetic defects. Unexpectedly, live-cell imaging revealed increased motility and frequent cyst fusion in SGEF-KD cysts. Restoring E-cadherin levels abolished cyst migration, while inhibition of matrix metalloproteinases markedly restored normal cyst volume and lumen architecture, identifying extracellular matrix remodeling as an additional contributor to the SGEF-deficient phenotype. Together, these findings identify SGEF as a key regulator of epithelial morphogenesis, coordinating junction integrity, actomyosin organization, lumen formation, and collective migration.

## Introduction

The epithelium is a monolayer of polarized cells that lines the outer surfaces of most mammalian organs and the inner surfaces of body cavities. These cells are connected by specialized junctional complexes that form a selectively permeable barrier separating the external environment from the organ’s interior (Rodriguez-Boulan & Macara, 2014). The establishment and maintenance of epithelial polarity and junction assembly are regulated by the Crumbs, Par, and Scribble protein complexes (Roignot et al., 2013). Beyond specifying apicobasal polarity, these protein complexes coordinate cell-cell adhesion, actomyosin organization, and interactions with the extracellular matrix to generate the mechanical forces required for epithelial morphogenesis (Buckley & St Johnston, 2022).

Among these critical regulators of polarity, the Scribble complex, composed of Scribble, Lethal giant larvae (Lgl), and Discs large (Dlg), plays a pivotal role in processes such as cell-cell adhesion, asymmetric cell division, vesicular trafficking, cell migration, and planar-cell polarity (Elsum et al., 2012). Because the Scribble complex lacks intrinsic catalytic activity, it is thought to function as a scaffolding platform that recruits downstream effectors which regulate these key cellular processes (Bonello & Peifer, 2019; Iden & Collard, 2008). In particular, Scribble-associated signaling pathways regulate members of the Rho family of small GTPases, which control cytoskeletal organization, junction dynamics, and cell migration (Etienne-Manneville & Hall, 2002). Dysregulation of these Rho GTPases, primarily driven by altered expression of their associated GEFs and GAPs, contributes to epithelial dysfunction and is implicated in a wide range of human cancers (Mack & Georgiou, 2014; Porter et al., 2016; Sahai & Marshall, 2002).

Our previous work in Madin-Darby Canine Kidney (MDCK) epithelial monolayers identified SGEF (ARHGEF26), a RhoG-specific GEF, as a component of the Scribble signaling network. SGEF, can simultaneously bind two members of the Scribble polarity complex, Scribble and Dlg1, and is recruited to the apical junctional complex in a Scribble-dependent manner, where it regulates adherens junction architecture, tight junction barrier integrity, and E-cadherin stability (Awadia et al., 2019). Stable knockdown of SGEF leads to increased junctional tension and transcriptionally mediated reductions in both ZO-1 and E-cadherin expression (Awadia et al., 2019; Rabino et al., 2024). Because E-cadherin and ZO-1 function not only as structural components of adherens and tight junctions but also as mechanosensitive platforms that couple cell-cell adhesion to the actomyosin cytoskeleton, disruption of their stability is expected to have profound consequences for epithelial tissue organization and morphogenesis.

While these findings show the critical role of SGEF in maintaining proper junction integrity, they were derived mostly from two-dimensional (2D) cell culture models. *In vivo*, epithelial cells grow in complex three-dimensional (3D) environments where cell-cell adhesion, cytoskeletal organization, tissue mechanics, and extracellular matrix remodeling must be precisely coordinated (Urzi et al., 2023). MDCK cysts provide a robust 3D model for studying epithelial polarization and morphogenesis, because they show similarities to key features of early organ development (Awadia et al., 2019; Bryant et al., 2010; Hirose et al., 2006). When cultured in Matrigel, single MDCK cells form spherical cysts composed of a polarized epithelial monolayer enclosing a central lumen. Lumen formation in this system depends on the coordinated establishment of apicobasal polarity, vesicle trafficking to the apical membrane initiation site (AMIS), maturation of epithelial junctions, and the generation of mechanical forces that allow the lumen to expand (Datta et al., 2011; Sigurbjornsdottir et al., 2014).

Our initial characterization of SGEF-deficient MDCK cysts demonstrated that silencing SGEF disrupts normal lumen formation, resulting in cysts containing multiple collapsed lumens (Awadia et al., 2019). However, these observations were primarily qualitative, leaving unresolved the mechanisms responsible for this striking phenotype.

Here, we combined quantitative morphometric analysis with long-term live-cell imaging to define the role of SGEF during epithelial morphogenesis in 3D MDCK cysts. We demonstrate that loss of SGEF disrupts lumen architecture, weakens epithelial junctions, alters actomyosin organization, promotes collective cyst migration and fusion, and enhances extracellular matrix remodeling. Mechanistically, reduced E-cadherin and ZO-1 expression downstream of increased β-catenin signaling contributes to impaired lumen expansion, while matrix metalloproteinase activity represents an additional pathway regulating cyst architecture. Together, our findings identify SGEF as a key signaling node that links the Scribble polarity complex to epithelial junction stability, cytoskeletal organization, and extracellular matrix remodeling, thereby coordinating the morphogenetic processes required to establish and maintain epithelial tissue architecture.

## Results

### SGEF regulates lumen number and morphology in MDCK cysts

In our previous study, we showed that SGEF regulates proper lumen formation in mature epithelial cysts without disrupting overall apicobasal polarity. CTRL cysts expressing a scrambled shRNA formed a single, spherical lumen, while SGEF KD cysts frequently developed multiple, collapsed lumens. These defects were rescued by lentiviral overexpression of mNeon-tagged human WT SGEF. Immunofluorescence (IF) also suggested that SGEF KD cysts had reduced E-cadherin levels compared to CTRL cysts (Awadia et al., 2019). However, these conclusions were based primarily on qualitative observations of single confocal z-planes. When the center z-plane of a cyst is imaged using confocal microscopy it is often difficult to conclusively determine whether the multiple luminal structures observed in SGEF KD cysts are independent lumens or a single multi-lobed lumen (Figure 1A).

**Figure 1.**
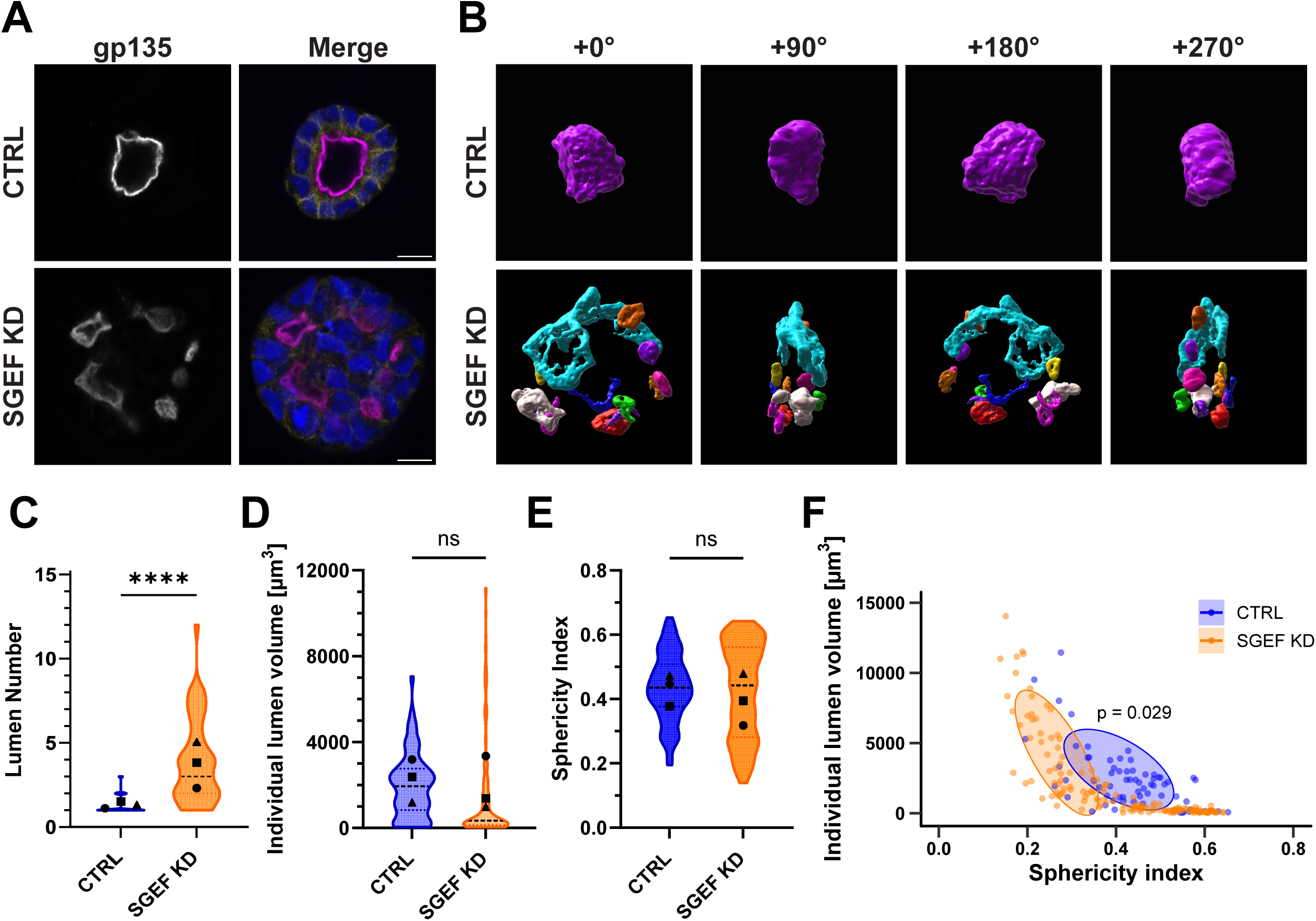
SGEF regulates lumen number and morphology in MDCK cysts. **(A)** Immunofluorescence of day 5 CTRL and SGEF KD cysts. Images show the central plane of each cyst. gp135 (magenta, cell apical surfaces/lumens), E-cadherin (yellow, adherens junctions), and Hoechst (blue, nuclei). Scale bars: 10 µm. **(B)** 3D Imaris reconstructions of lumens from (A). “+x⁰” indicates rotation along the vertical axis. **(C)** Lumen number per cyst. **(D)** Individual lumen volume. **(E)** Lumen sphericity. **(F)** Relationship between lumen volume and sphericity. Each point represents one lumen. Ellipses surround 60% of primary luminal structures (> 1000 µm^3^). PERMANOVA P value is shown on the graph.For (C), n=3 independent experiments (>15 cysts/condition/experiment). For (D-F), n=3 (>15 lumens/condition/experiment). Violin plots show data distribution; points represent experiment means. Center line, median; bounds, interquartile range (IQR). ns=not significant. ****P<0.0001. See Methods for quantitative and statistical details.

To better define the role of SGEF in lumen formation, we developed new image segmentation pipelines in Fiji/ImageJ to quantitatively analyze different morphometric parameters of 3D cysts and their internal lumens. These included lumen number and shape, cyst volume, and cell number, among others (see Methods). We first quantified the number of lumens per cyst in CTRL *vs.* SGEF KD cells. To better visualize luminal structures, we used Imaris to segment and render mature CTRL and SGEF KD cysts in 3D, which resolved multiple distinct lumens in SGEF KD cysts, as predicted (Fig. 1B, Supp. Video 1A, B). Quantification over multiple experiments showed a significant, ∼4-fold increase in lumen number in SGEF KD cysts over CTRL cysts, with an average of 1.30 +/- 0.08 lumens in CTRL cysts and 3.87 +/- 0.36 lumens in SGEF KD cysts (Fig. 1C).

SGEF KD cysts often displayed elongated and flattened lumens, which in some extreme cases, appeared to be completely devoid of fluid or “closed.” To better characterize this phenotype, we quantified both the volume and shape of individual lumens. Lumen shape was assessed using sphericity, which is an index ranging from 0 to 1, where 0 describes an object that is completely flat and 1 describes a perfect sphere (Wadell, 1932). Our results show that both individual lumen volume and sphericity appeared to be lower in SGEF KD cysts, but the results were not statistically significant. (Fig. 1D, E). The shape of the violin plots, especially for individual lumen volume, shows that SGEF KD cysts have a markedly different distribution than CTRL cysts, with a broader range of lumen volumes and an enrichment of very small lumens. This distribution suggests the presence of distinct lumen populations with different morphological characteristics. Plotting each lumen’s volume with its corresponding sphericity index revealed these distinct subpopulations of lumens, including small, highly spherical secondary lumens and larger primary lumens that differed in morphology between experimental conditions. CTRL lumens had a relatively homogeneous distribution, clustering within a narrow range of volumes and maintaining a roughly spherical shape, as expected (Fig. 1F, blue). In contrast, SGEF KD lumens showed substantial heterogeneity, spanning a wide range of volumes and sphericities (Fig. 1F, orange). Interestingly, SGEF KD lumens with sphericity index values comparable to CTRL lumens (0.4-0.6) were generally much smaller in volume (500-2000 µm^3^ vs. 500-4900 µm^3^ in CTRL). As lumen volume increased, SGEF KD lumens became progressively less spherical, reaching sphericity values as low as 0.18 at volumes between 2500 and 14000 µm^3^. These findings suggest that, in the absence of SGEF, lumens are unable to maintain a spherical morphology during lumen expansion (Fig. 1F).

### SGEF regulates cell-cell junction protein levels in 3D epithelial cysts

Disruption of the Scribble/SGEF/Dlg1 ternary complex by silencing SGEF expression in 2D culture reduces the expression of the junctional proteins E-cadherin and ZO-1 (Awadia et al., 2019; Rabino et al., 2024). To assess whether SGEF similarly regulates junctional organization in a 3D environment, we quantified the intensity of multiple junctional proteins in mature MDCK cysts using IF and western blot (WB).

For IF measurements, we developed an image analysis pipeline that measures the total fluorescence intensity of each junctional marker throughout the entire cyst and normalizes this value to the number of cells per cyst (see Methods). Using this approach, we found that SGEF KD cysts had significantly reduced E-cadherin (∼5-fold) and ZO-1 (∼2-fold) fluorescence compared with CTRL cysts, consistent with our previous findings in monolayers. Re-expression of WT SGEF restored E-cadherin expression to levels that were 1.7-fold higher than CTRL, whereas ZO-1 expression was only partially rescued (Fig. 2A, C-D).

**Figure 2.**
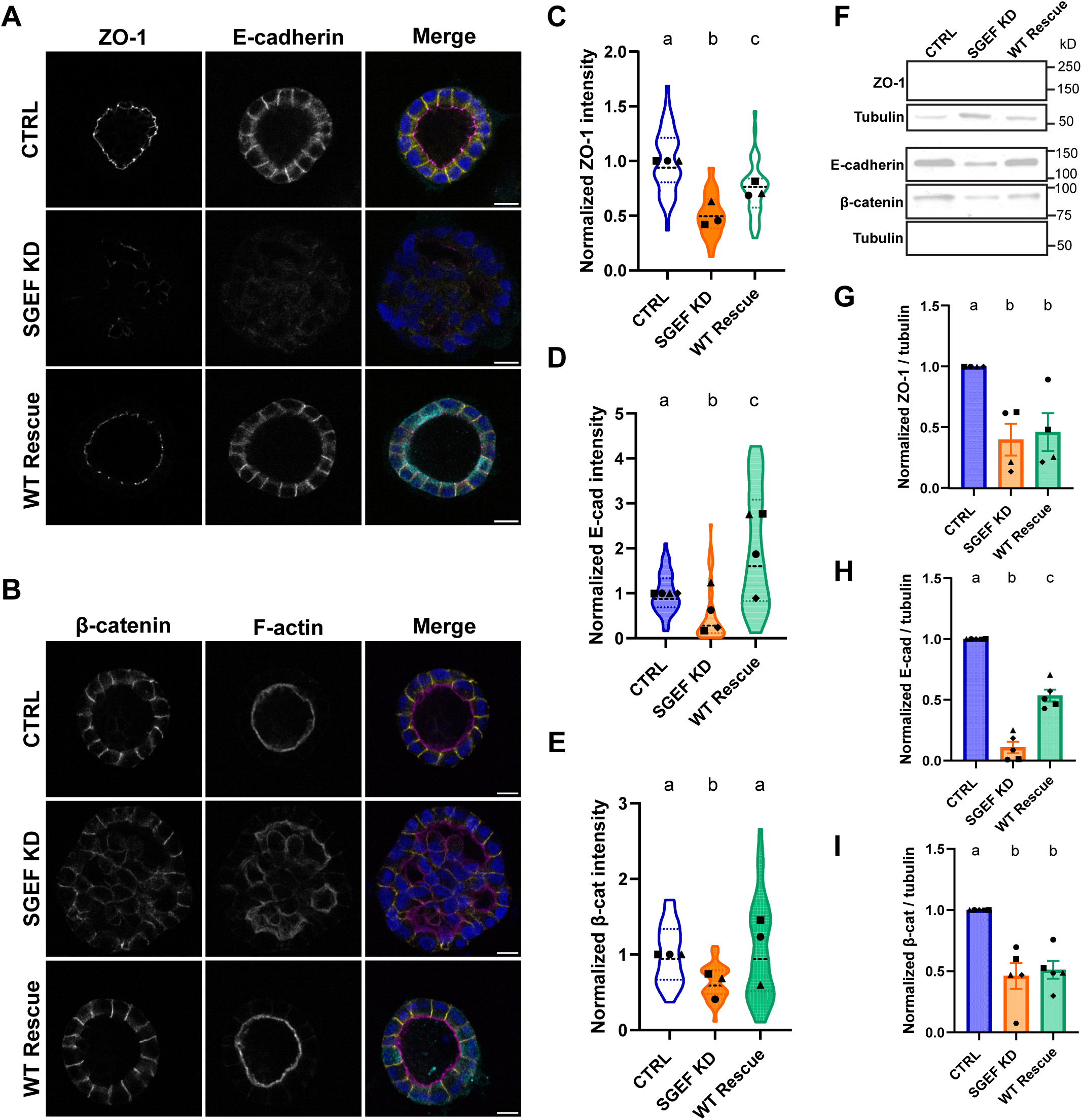
SGEF regulates cell-cell junction protein levels in 3D epithelial cysts. **(A)** Immunofluorescence of CTRL, SGEF KD, and WT Rescue cysts stained for ZO-1 (magenta), E-cadherin (yellow), mNeon (cyan), and Hoechst (blue). WT Rescue expresses mNeon-tagged WT SGEF in SGEF KD cells. Scale bars: 10 µm. **(B)** Immunofluorescence of β-catenin (yellow), F-actin (magenta), mNeon (cyan), and Hoechst (blue). Scale bars: 10 µm. **(C-E)** Quantification of ZO-1, E-cadherin, and β-catenin fluorescence intensity, normalized to CTRL. Intensity was quantified as the summed integrated density across all z-planes and normalized to cell number. **(F)** Representative western blots of ZO-1, E-cadherin, and β-catenin from total cyst lysates. Tubulin was used as a loading control. **(G-I)** Quantification of ZO-1, E-cadherin, and β-catenin protein levels normalized to CTRL. All cysts were cultured for 7 days in experiments contributing to this figure. For (C), n=3 independent experiments (>15 cysts/condition/experiment); (D), n=4 (>15 cysts/condition/experiment); (E), n=5 (>15 cysts/condition/experiment); (G-I), n≥4. Violin plots show data distribution; symbols represent the means of each individual experiment. Center line, median; bounds, IQR. Conditions not sharing a letter are significantly different (P<0.05, Dunn’s multiple comparisons test).

Our previous results also showed that, in 2D, β-catenin protein levels were unchanged following SGEF KD, despite displaying a more diffuse junctional localization (Awadia et al., 2019). We therefore investigated β-catenin expression in 3D cysts. Interestingly, quantification of IF images revealed that the average β-catenin levels were lower in SGEF KD cysts, when compared to CTRL and Rescue WT conditions (Fig. 2B, E). This discrepancy may reflect differences in gene expression and protein regulation between 2D and 3D culture systems, which are known to have distinct transcriptomes in many cases (Liu et al., 2022; Storch et al., 2010)

To validate the IF-based quantification and distinguish changes in protein abundance from changes in subcellular localization, we analyzed protein expression by western blotting of lysates prepared from mature 3D cysts. Western blot measurements showed greater experimental variability, likely reflecting the technical challenges associated with recovering protein from cysts embedded in Matrigel, which contains a high concentration of laminin. Despite this variability, the overall trends for E-cadherin, ZO-1, and β-catenin resembled the IF measurements (Fig. 2F-I). Collectively, these findings demonstrate that SGEF is required to maintain AJ and TJ integrity in 3D epithelial cysts.

### SGEF regulates TJ architecture and actomyosin distribution in 3D cysts

Our previous work showed that silencing SGEF in 2D MDCK monolayers increased basal actin stress fiber formation and altered tight junction architecture, resulting in straighter junctions accompanied by a striking redistribution of non-muscle myosin IIB into junctional arrays (Awadia et al., 2019). To determine whether SGEF similarly regulates the actomyosin network in 3D epithelial cysts, we examined F-actin, tight junction morphology, and myosin IIB localization.

Surprisingly, F-actin staining showed a predominantly apical distribution in both CTRL and SGEF KD cysts, with minimal recruitment to apical junctions in SGEF KD (Fig. 2B, Supp. Fig. 1). This contrasts our observations of basal stress fibers in 2D monolayers and may reflect the compliance of the Matrigel in 3D cysts, as opposed to the stiffness associated with cells plated on glass.

Despite the similar F-actin distribution between conditions, tight junction organization remained altered in SGEF KD cysts. ZO-1 staining showed junctions that appeared more linear compared with the curvilinear morphology observed in CTRL cysts (Fig. 3A, Supp. Video 2A). Re-expression of WT SGEF restored normal junction morphology (Fig. 3A, Supp. Video 2C). To quantify the degree of linearity displayed by tight junctions, we measured their zig-zag index, which is defined as the ratio of the length of a freehand line traced along the shape of a tight junction from cell vertex to vertex to that of a straight line drawn from cell vertex to vertex (Tokuda et al., 2014). Our results confirmed a reduction in the tortuosity of tight junctions in SGEF KD cysts, which was restored upon re-expression of WT SGEF (Fig. 3B).

**Figure 3.**
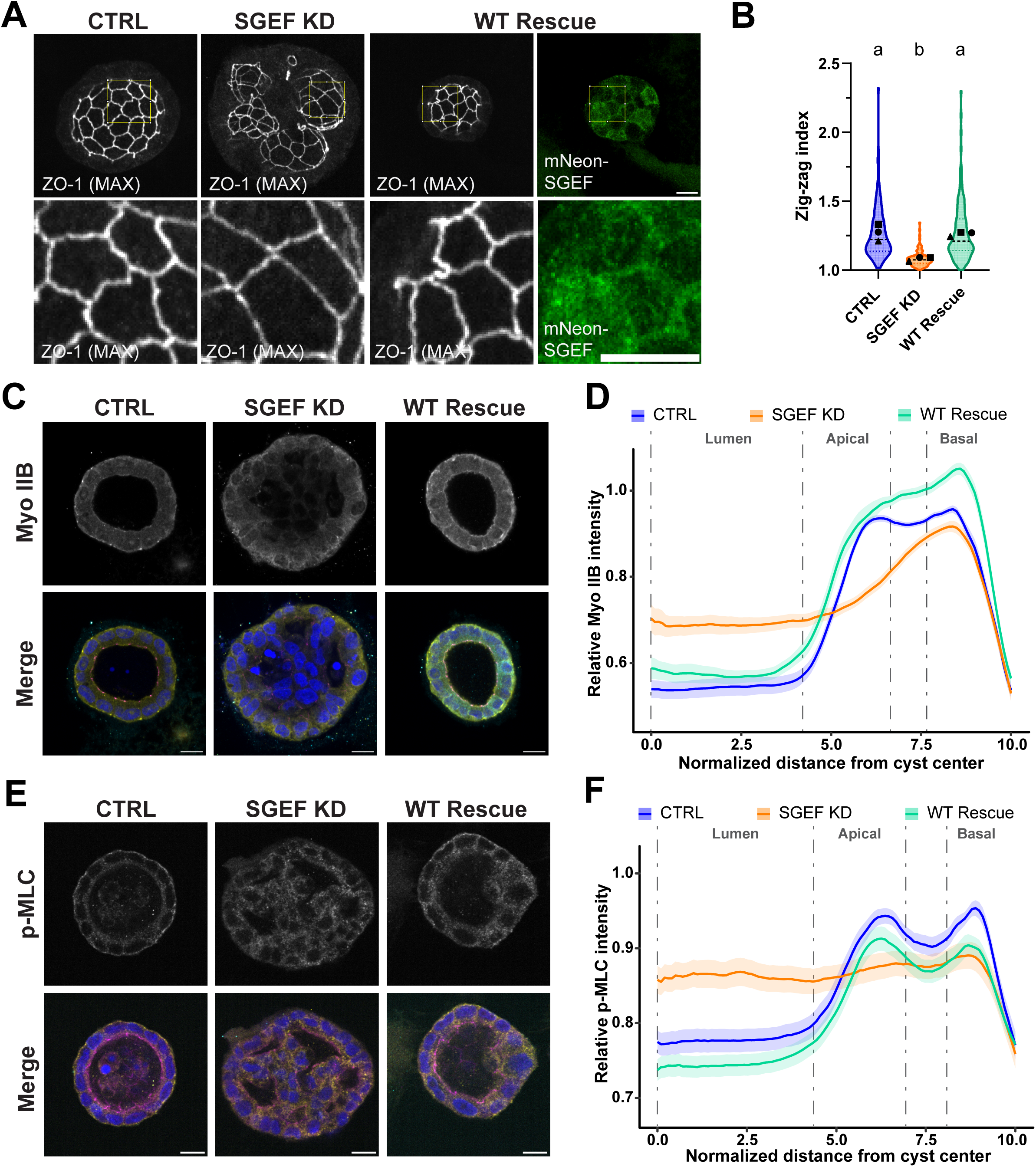
SGEF coordinates TJ architecture and actomyosin organization in 3D cysts. **(A)** ZO-1 immunofluorescence in CTRL, SGEF KD, and WT Rescue cysts. Maximum intensity projections were generated from 10-20 consecutive 0.2 µm z-slices beginning at the most basal, visible region of the junctional network. Zoomed region in the bottom panel is indicated with a yellow square. Scale bars: 10 µm. **(B)** Quantification of tight junction “zig-zag index” measured from projections similar to those shown in A (n=3 independent experiments; >5 random junctions/cyst from 10 cysts/condition/experiment). **(C)** Non-muscle myosin IIB (yellow), ZO-1 (magenta), mNeon (cyan), and Hoechst (blue) staining. Scale bars: 10 µm. **(D)** Radial intensity profiles of non-muscle myosin IIB from cyst center to periphery (n=3; >10 cysts/condition/experiment). **(E)** p-MLC (yellow), ZO-1 (magenta), mNeon (cyan), and Hoechst (blue) staining. Scale bars: 10 µm. **(F)** Radial p-MLC intensity profiles (n=2; >10 cysts/condition/experiment). All cysts were cultured for 7 days in experiments contributing to this figure.

Non-muscle myosin IIB is known to be differentially localized in 3D cysts compared to monolayers (Cerruti et al., 2013; Yu et al., 2008). As expected, it was enriched along the basal surfaces, with some enrichment at the apical membrane in both CTRL and WT Rescue cysts. In contrast, SGEF KD cysts showed a more diffuse myosin IIB distribution throughout the epithelium, accompanied by increased localization at internal cell-cell contacts and highly diffuse enrichment at the basal surface (Fig. 3C; Supp. Fig. 2A, B). To quantify these changes, we adapted a radial intensity analysis pipeline previously developed in our lab to measure intensity at invadopodia (Kreider-Letterman et al., 2023). This semi-automatic script measures the average intensity of a particular protein in relation to the center of the cyst (see Methods for details). Our results confirmed the redistribution of myosin IIB in SGEF KD cysts, from apical and basal localization in CTRL cysts, to a more diffuse pattern with a less pronounced basal enrichment (Fig. 3D).

To determine whether active myosin followed a similar pattern, we also conducted immunostaining for phosphorylated myosin light chain (p-MLC). Consistent with the myosin IIB results, p-MLC displayed apical and basal enrichment in CTRL and Rescue WT cysts, and a broader and more diffuse distribution in SGEF KD cysts (Fig. 3E; Supp. Fig. 2C, D). However, unlike myosin IIB localization, SGEF KD cysts did not have asymmetrical enrichment of active myosin at any location within the cyst (Fig. 3F).

This data demonstrates that loss of SGEF alters both the architecture of the junctions and the organization of the actomyosin network in 3D epithelial cysts. The reduced junctional curvature observed in SGEF KD cysts suggests elevated junctional tension, similar to our previous observations in monolayers. Although myosin IIB did not assemble into the junctional arrays observed in 2D, its redistribution toward cell-cell contacts may contribute to increased contractility between cells. These findings support the idea that SGEF plays a role in regulating epithelial contractility and junctional mechanics.

### SGEF governs lumen size and architecture through β-catenin-dependent regulation of E-cadherin and ZO-1

We have recently demonstrated that SGEF KD destabilizes E-cadherin at cell-cell junctions, promoting its internalization via clathrin-mediated endocytosis (Rabino et al., 2024). This process disrupts the E-cadherin/catenin complex, releasing β-catenin to act as a co-activator of Slug, a transcriptional repressor of E-cadherin, leading to its downregulation (Rabino et al., 2024). We also observed that SGEF KD decreases transcription of the tight junction protein ZO-1 through a β-catenin-dependent mechanism, although the underlying molecular pathway remains unknown.

To determine the contributions of E-cadherin and ZO-1 downregulation to lumen morphogenesis, we treated SGEF KD cysts with the β-catenin signaling inhibitor iCRT3. We previously demonstrated that iCRT3 restores E-cadherin and ZO-1 expression, as well as tight junction integrity in 2D monolayers (Rabino et al., 2024). iCRT3 functions by directly binding β-catenin and preventing its interaction with transcription factors but has no effect on its interaction with E-cadherin (Gonsalves et al., 2011). Consistent with our findings in 2D cultures, iCTR3 treatment successfully restored ZO-1 and E-cadherin expression in 3D cysts (Fig. 4A-C).

**Figure 4.**
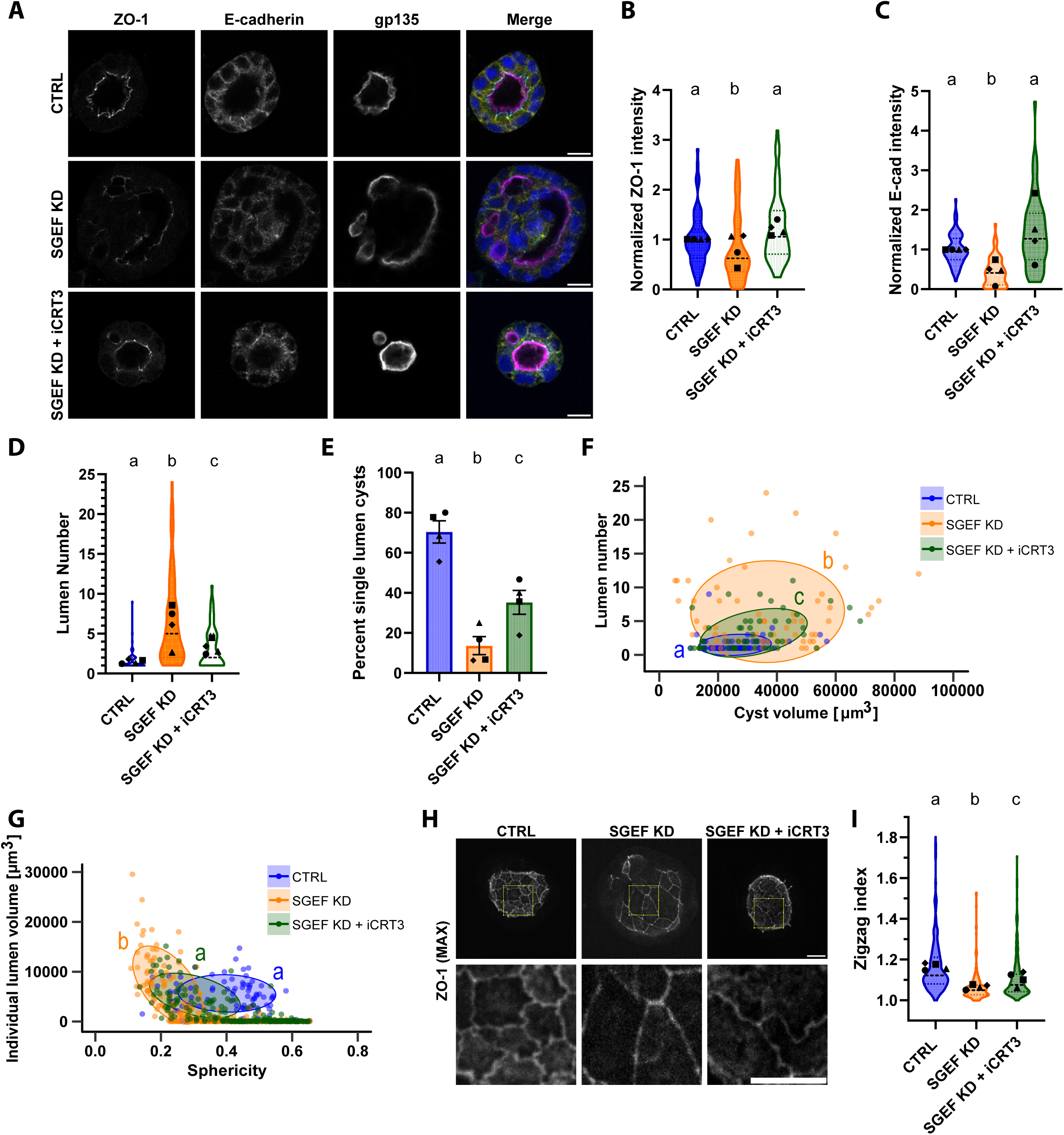
SGEF governs lumen size and architecture through β-catenin-dependent regulation of E-cadherin and ZO-1. **(A)** Immunofluorescence of CTRL, SGEF KD, and SGEF KD + iCRT3 cysts stained for ZO-1 (cyan), E-cadherin (yellow), gp135 (magenta), and Hoechst (blue). Scale bars: 10 µm. **(B-C)** ZO-1 and E-cadherin intensity, normalized to CTRL. **(D)** Lumen number per cyst. **(E)** Percentage of cysts with a single lumen. **(F)** Relationship between lumen number and cyst volume. Each point represents one cyst, and ellipses encompass 60% of each condition. Letters indicate statistical groupings. **(G)** Individual lumen volume versus sphericity. Each point represents one lumen. Ellipses encompass 60% of lumens >1000 µm^3^ in each condition. **(H)** ZO-1 maximum intensity projections, with regions of interest (ROIs) enhanced below. Scale bars: 10 µm. **(I)** Tight junction zig-zag index. Cysts were cultured for 7 days in all experiments contributing to this figure. For (B-F), n=4 independent experiments (>15 cysts/condition/experiment). For (G), n=4 (1-25 lumens/cyst from >15 cysts/condition/experiment). For (I), n=4 (>5 random junctions/cyst from 10 cysts/condition/experiment). Violin plots show data distribution; symbols represent the means of each individual experiment. Center line, median; bounds, IQR. Conditions not sharing a letter are significantly different (P<0.05 Dunn’s test or PERMANOVA).

Importantly, restoring E-cadherin and ZO-1 expression partially rescued the lumen morphogenesis defects caused by SGEF depletion (Fig. 4A, D). The average number of lumens per cyst decreased from 6.46 +/- 0.72 lumens per cyst in SGEF KD to 3.28 +/- 0.33 lumens per cyst following iCRT3 treatment, compared with 1.53 ± 0.14 lumens per cyst in control cysts. Similarly, iCRT3 treatment significantly increased the percentage of cysts containing a single lumen from 13.65 +/- 4.49% in SGEF KD to 35.28 +/- 5.95% after iCRT3 treatment. However, this rescue was partial, because the percentage of single lumen cysts remained substantially lower than in CTRL cysts (70.44 +/- 5.56%) (Fig. 4E). Analysis of the relationship between cyst volume and lumen number also suggested a partial rescue following iCRT3 treatment, although this trend did not reach statistical significance, most likely because of the considerable variability among SGEF KD cysts (Fig. 4F). In contrast, the relationship between individual lumen volume and sphericity was restored to CTRL levels (Fig. 4G).

In addition to improving lumen number and morphology, iCRT3 substantially restored tight junction organization. Whereas SGEF KD cysts exhibited a disorganized tight junction network, iCRT3-treated cysts displayed the characteristic cage-like ZO-1 organization surrounding the primary lumen, closely resembling CTRL cysts (Supp. Video 3). Furthermore, iCRT3 restored the curvilinear morphology of tight junction (Fig. 4H). This was confirmed quantitatively by measuring the zig-zag index, which showed a partial rescue following iCRT3 treatment (Fig. 4I).

Together, our results demonstrate that β-catenin-dependent downregulation of E-cadherin and ZO-1 contributes to the collapsed, multi-lumen phenotype observed in SGEF KD cysts. However, restoring E-cadherin and ZO-1 expression in the absence of SGEF was insufficient to fully rescue normal lumen morphogenesis, suggesting that SGEF performs additional functions that are critical for epithelial cyst development.

### Cyst volume, cell number, and proliferation are influenced by SGEF

During analysis of the results described above (see Figs. 1A. 2A-B, 3C-D, Fig. 4A), it became apparent that SGEF KD cysts tended to be larger than both CTRL and WT Rescue cysts cultured under identical conditions. Quantification confirmed these observations. Mature SGEF KD cysts showed increased volume and were comprised of more cells, averaging 74,123 µm^3^ +/- 3,482 µm^3^ and 96.76 +/- 3.64 nuclei per cyst, compared with 43,878 µm^3^ +/- 3,553 µm^3^ and 63.64 +/- 3.02 nuclei in CTRL, along with 41,305 µm^3^ +/- 2,642 µm^3^ and 68.01 +/- 3.43 nuclei in WT Rescue cysts (Fig. 5A-C). The relationship between cell number and cyst volume showed similar slopes among CTRL, SGEF KD, and WT Rescue cysts, suggesting that larger cyst volumes were associated with a proportional increase in cell number (Fig. 5D).

**Figure 5.**
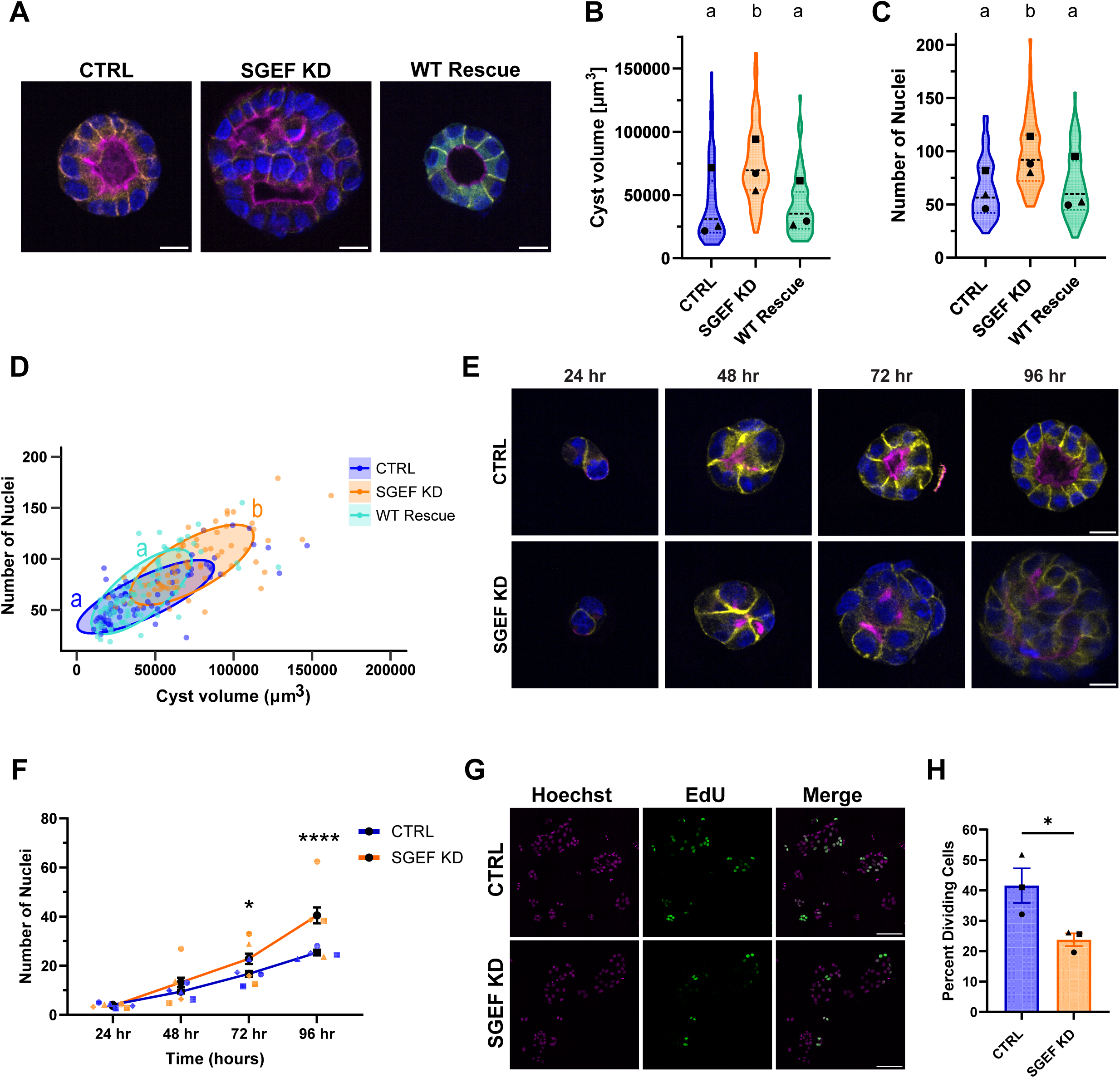
SGEF silencing affects cyst volume, cell number, and cell proliferation. **(A)** Immunofluorescence of day 7 CTRL, SGEF KD, and WT Rescue cysts stained for E-cadherin (yellow), F-actin (magenta), mNeon (cyan), and Hoechst (blue). Scale bars: 10 µm. **(B)** Cyst volume. **(C)** Nuclei per cyst. **(D)** Relationship between cyst volume and nuclei number. Each point represents one cyst; ellipses encompass 60% of each condition, and letters indicate statistical groupings. **(E)** Time course of cyst development (24-96 h) stained for E-cadherin (yellow), gp135 (magenta), and nuclei (blue). **(F)** Nuclei number over time. Black symbols indicate the mean +/- SEM of all experiments combined. Blue (CTRL) and orange (SGEF KD) symbols indicate the mean of each individual experiment. **(G)** Representative images of EdU incorporation in 2D MDCK cultures after 3 days. Hoechst (magenta), EdU (green). Scale bars: 100 µm. **(H)** Percentage of EdU-positive cells per field of view (FOV). For (B-D), n=3 independent experiments (>15 cysts/condition/experiment). For (F), n=4 (>15 cysts/time point/condition/experiment). For (H), n=3 (5 FOVs/condition/experiment). Violin plots show data distribution; symbols represent the means of each individual experiment. Center line, median; bounds, IQR. Conditions not sharing a letter are significantly different (P<0.05; Dunn’s test or PERMANOVA). *P<0.05. ****P<0.0001. See Methods for statistical analysis.

To investigate when differences in cell number first occurred, we fixed cysts at multiple developmental time points between 24 and 96 hours after seeding and quantified the number of nuclei per cyst (Fig. 5E, F). We found that divergence in cell number was apparent as early as 48 hours, with SGEF KD cysts averaging 13.24 +/- 1.88 nuclei, compared with 9.45 +/- 0.75 nuclei in CTRL cysts. These differences became statistically significant by 72 hours, where SGEF KD cysts contained an average of 22.85 +/- 2.07 nuclei compared with 16.69 +/- 1.17 nuclei in CTRL cysts (Fig. 5E, F). These findings indicate that loss of SGEF leads to an early increase in cell number during cyst development.

To determine whether increased cell number resulted from a higher proliferation rate, we performed an EdU incorporation assay in 2D monolayers cultured for 72 hours. Surprisingly, SGEF KD cells showed a significantly lower proportion of EdU-positive cells than CTRL cells, with 23.78 +/- 2.09% proliferating cells compared to 41.61 +/- 5.67% in CTRL cells (Fig. 5G, H). Together, these results suggest that the increased cyst volume and cell number observed following SGEF silencing could not be attributed to increased proliferation alone, suggesting that alternative mechanisms contribute to large SGEF KD cysts.

### Silencing SGEF stimulates cyst motility and fusion

To investigate the mechanisms underlying the increased volume of SGEF KD cysts over time, we performed live imaging of developing cysts over 84 hours, beginning 12 hours after seeding. CTRL and SGEF KD cells stably expressing an mScarlet-CAAX membrane marker were used to visualize cyst morphology and behavior during development. Drastic behavioral differences were immediately apparent between conditions (Fig. 6A). Although cyst rotation and cell division were observed in both CTRL and SGEF KD cysts, SGEF KD cysts showed increased motility, frequently extending one or a small group of cells ahead of the main cyst body while remaining physically connected (Fig. 6A, B, Supp. Video 4). Quantitative analysis confirmed that SGEF KD cysts traveled a greater total distance (437.54 µm +/- 38.84 µm) than CTRL cysts (318.85 µm +/- 23.35 µm) (Fig. 6C). SGEF KD cysts also migrated at higher average velocities (5.73 µm/h +/- 0.38 µm/h) and with increased motility persistence (0.11 +/- 0.01) compared to CTRL cysts (4.14 µm/h +/- 0.17 µm/hr and 0.09 +/- 0.01) (Fig. 6D, E).

**Figure 6.**
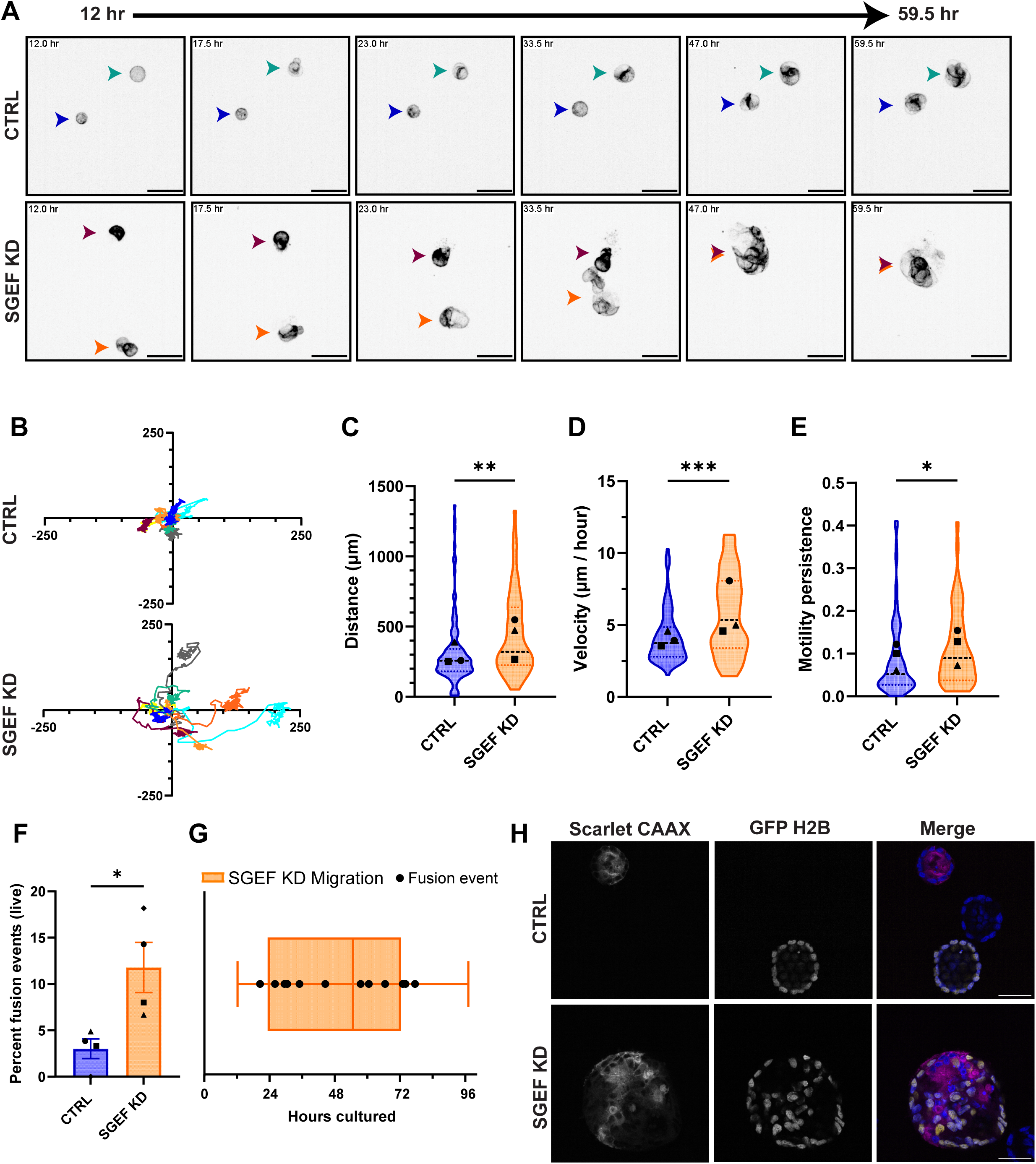
Silencing SGEF promotes cyst motility and fusion. **(A)** Representative live imaging of CTRL and SGEF KD cysts expressing Scarlet-CAAX (plasma membrane marker), shown from 12-59.5 h after seeding. Scale bars: 40 µm. **(B)** Representative migration tracks of 10 individual cysts per condition, normalized to their starting positions. **(C)** Total migration distance per cyst. **(D)** Mean migration velocity per cyst (µm/h). **(E)** Average motility persistence per cyst (net displacement/total path length). **(F)** Frequency of live cyst fusion events. **(G)** Timing of migration and fusion events in SGEF KD cysts. The box represents average time of cyst migration start and end. Whiskers represent the range of observed cyst migration time, while the vertical line represents the median of that range. Each point represents the timing of one fusion event between two cysts. **(H)** Scarlet-CAAX and GFP-H2B expressing cells were co-plated in Matrigel to monitor and quantify cyst fusion. Representative images of CTRL and SGEF KD mixed labeled populations as cysts. The bottom panel shows a fusion event in SGEF KD cysts, marked by a cyst expressing both markers in different cells within the cyst. Scale bars: 30 µm. For (C-F), n=3 (>10 cysts/condition/experiment). For (G), n=3 (∼4 fusion events/experiment). Violin plots show data distribution; symbols represent experiment means. Center line, median; bounds, IQR. *P<0.05. **P<0.01. ***P<0.001. See Methods for statistical analysis.

During these migratory events, SGEF KD cysts frequently contacted and fused with neighboring cysts, providing a potential explanation for their increased volume in fixed samples (Fig. 6A, B, Supp. Video 4). Analysis of live-imaging datasets revealed that an average of 11.78 +/- 2.70% of SGEF KD cysts underwent fusion events, compared to only 3.01 +/- 1.06% of CTRL cysts (Fig. 6F). To further characterize the relationship between migration and fusion, we quantified the timing of both processes throughout cyst development. SGEF KD cysts showed the highest levels of migration between 24 and 72 hours after seeding, with fusion events occurring over a similar time frame (Fig. 6G). Notably, this period coincided with the onset of divergence in cell number between CTRL and SGEF KD cysts.

As a complementary approach to quantify cyst fusion, we designed an assay in which two populations of cells were co-seeded for each condition: one stably expressing mScarlet-CAAX and the other expressing nuclear GFP-H2B. Cultures were fixed after seven days and examined for dual-labeled cysts (Fig. 6H). Because mature cysts typically arise from a single cell, cysts containing both mScarlet and GFP fluorescence were scored as cysts that had undergone fusion events. The percentage of dual-labeled cysts closely matched the values obtained from live imaging (6.76 +/- 0.86% in CTRL and 17.01 +/- 0.39% in SGEF KD), independently confirming that fusion occurs more frequently in SGEF KD cysts than in CTRL cysts (Supp. Fig. 3A).

The observation of cyst fusion in SGEF KD cells motivated us to investigate whether cyst fusion could account for the multi-lumen phenotype. If fusion were the primary cause, then smaller, non-fused cysts would be expected to contain a higher proportion of single lumen. To test this possibility, we examined the relationship between cyst volume and lumen number in CTRL and SGEF KD cysts. Our results show that, even though most SGEF KD cysts were substantially larger than CTRL cysts, when cysts of similar volumes where compared, SGEF KD cysts consistently contained more lumens (Supp. Fig. 3B, C, dashed lines). These findings indicate that the abnormal lumen morphology observed following SGEF knockdown cannot be attributed to cyst fusion alone.

### Restoring E-cadherin in SGEF KD cysts is sufficient to rescue migration

Loss of E-cadherin is a hallmark of EMT and is frequently associated with increased migratory behavior in both developmental and cancer contexts (Batlle et al., 2000; Behrens et al., 1993; Cano et al., 2000; Perl et al., 1998; Vleminckx et al., 1991). Because SGEF depletion reduces E-cadherin expression, we investigated whether E-cadherin loss contributes to the migratory phenotype observed in SGEF KD cysts. As previously stated, our lab demonstrated that E-cadherin downregulation in SGEF KD cells is mediated by increased expression of the transcriptional repressor Slug. Stable knockdown of Slug in SGEF KD cells (SGEF/Slug dKD) restored E-cadherin expression to levels comparable to CTRL cells without rescuing ZO-1 expression (Rabino et al., 2024). To assess the functional consequences of E-cadherin restoration during cyst development, we generated SGEF/Slug dKD cells stably expressing mScarlet-CAAX and conducted live imaging from 12 to 96 hours after seeding.

Restoration of E-cadherin drastically reduced the migratory behavior observed in SGEF KD cysts (Fig. 7A, B, Supp. Video. 5). Quantitative analysis showed that the total distance traveled by SGEF/Slug KD cysts, on average, was significantly reduced (321.28 µm +/- 38.08 µm) compared to SGEF KD cysts (605.88 µm +/- 73.30 µm), and approached CTRL levels with (261.08 µm +/- 15.69 µm) (Fig. 7C). Also, SGEF/Slug dKD cysts migrated at lower velocities (4.09 µm/h +/- 0.53 µm/h) and demonstrated reduced directional persistence (0.09 +/ 0.02) relative to SGEF KD cysts (7.81 µm/h +/- 0.55 µm/h and 0.16 +/- 0.02) (Fig. 7D, E), with values that were not statistically different from CTRL cysts. In addition to abolishing migratory behavior, Slug KD also markedly reduced fusion events (0.89 +/- 0.89%) to levels lower than both SGEF KD (21.91 +/- 8.57%) and CTRL (7.18 +/- 3.55%) (Fig. 7F). These findings indicate that E-cadherin loss is a major contributor to the enhanced motility and fusion caused by knockdown of SGEF, and its rescue is sufficient to prevent a migratory phenotype.

**Figure 7.**
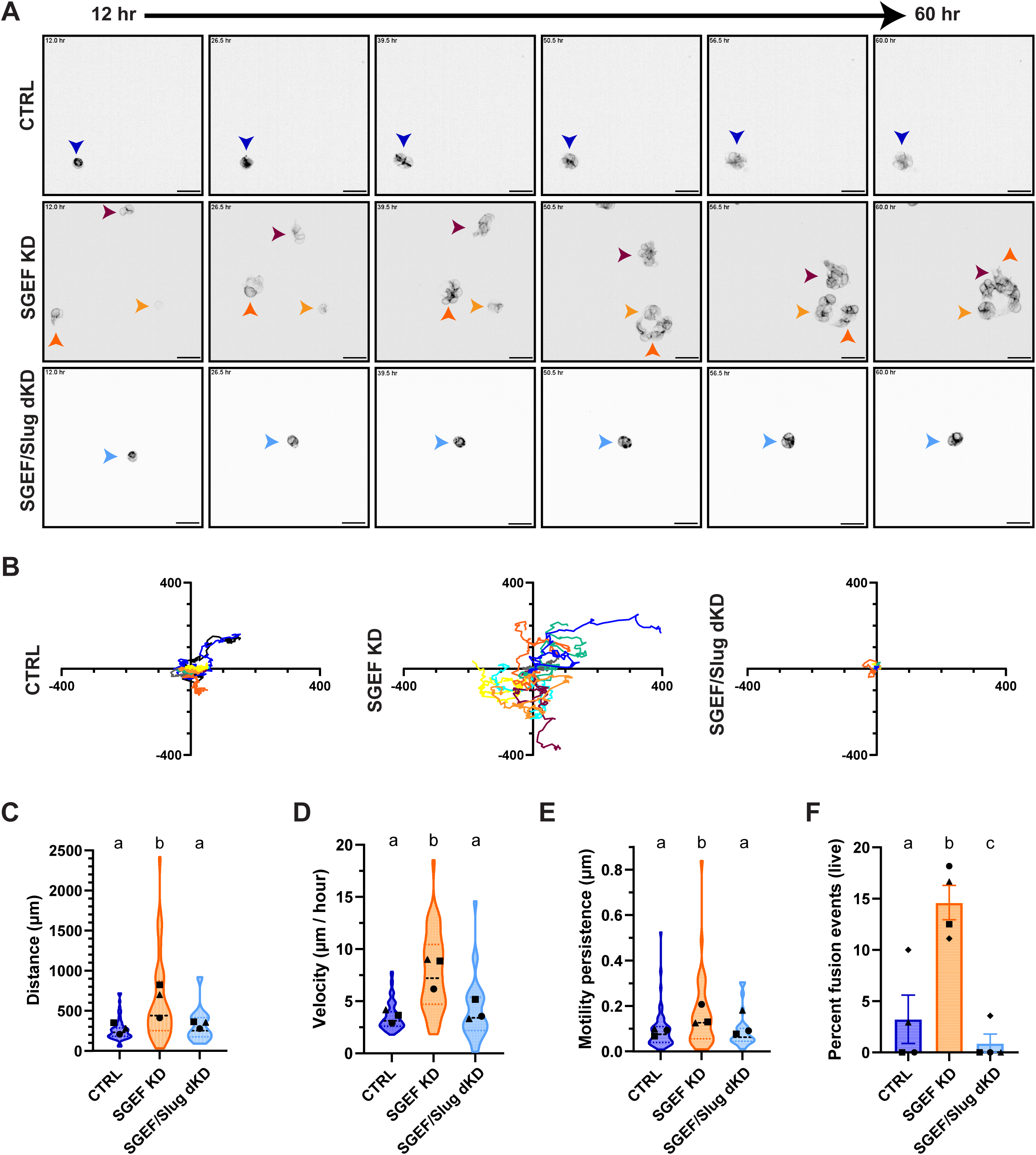
SGEF KD cyst migration is mediated by E-cadherin expression. **(A)** Live imaging of CTRL, SGEF KD, and SGEF/Slug KD cysts expressing Scarlet-CAAX. Representative frames from 12-60 h after seeding are shown. Scale bars: 40 µm. **(B)** Representative migration tracks of 10 individual cysts per conditions imaged over 84 h. **(C)** Total migration distance. **(D)** Mean migration velocity (µm/hour). **(E)** Motility persistence. **(F)** Frequency of live cyst fusion events. For (C-F), n=3 (>10 cysts/condition/experiment). Violin plots show data distribution; symbols represent experiment means. Center line, median; bounds, IQR. Conditions not sharing a letter are significantly different (P<0.05, Dunn’s test).

### Metalloproteinase activity influences cyst volume and lumen phenotype in SGEF KD cysts

Enhanced cell migration and invasion in 3D is frequently associated with increased matrix metalloproteinase (MMP) expression and/or activity, which promotes extracellular matrix (ECM) remodeling through proteolytic degradation of matrix components, including collagens and laminins (Egeblad & Werb, 2002). We therefore examined whether MMP activity contributed to the migratory and morphogenetic defects observed following SGEF KD.

To inhibit MMP activity, CTRL and SGEF KD cysts were cultured in the presence or absence of GM6001, a broad-spectrum MMP inhibitor (Fig. 8A). GM6001 treatment had no detectable effect on CTRL cyst morphology, with most treated cysts displaying a single central lumen (Fig. 8A, E). In contrast, MMP inhibition produced two striking effects in SGEF KD cysts. First, GM6001 treatment reduced SGEF KD cyst volume of by nearly 50%, from 78,827 µm^3^ +/- 4,785 µm^3^ to 44,695 µm^3^ +/- 3,028 µm^3^ (Fig. 8B). Because MMP inhibition is not expected to directly affect proliferation, this reduction in cyst size likely reflects decreased cyst migration and fusion during development. These findings suggest that the increased motility observed in the absence of SGEF is MMP-dependent. Second, GM6001 treatment unexpectedly rescued the multi-lumen phenotype in SGEF KD cysts (Fig. 8A, C, E). The average number of lumens per SGEF KD cyst decreased from 5.21 +/- 0.58 to 2.59 +/- 0.27 following GM6001 treatment (Fig. 8C). Consistent with this observation, plotting lumen number against cyst volume revealed a partial but significant rescue, characterized by a prominent cluster of smaller cysts containing a single lumen (Fig. 8D). Remarkably, MMP inhibition restored the SGEF KD phenotype more effectively than inhibition of β-catenin signaling with iCRT3 (compare Figs. 4D and 4F with 8C and 8D). To further assess this effect, we quantified the proportion of single-lumen cysts across all GM6001 experiments. In untreated SGEF KD cultures, only 16.81 +/- 10.34% of cysts were single lumen, whereas GM6001 treatment increased this percentage to 42.41 +/- 6.50%. This rescue was comparable to, and trended greater than, that achieved with iCRT3 treatment (35.28 +/- 5.95%) (Fig. 4E), suggesting that MMP activity contributes at least as strongly as β-catenin signaling in determining lumen number during cyst morphogenesis.

**Figure 8.**
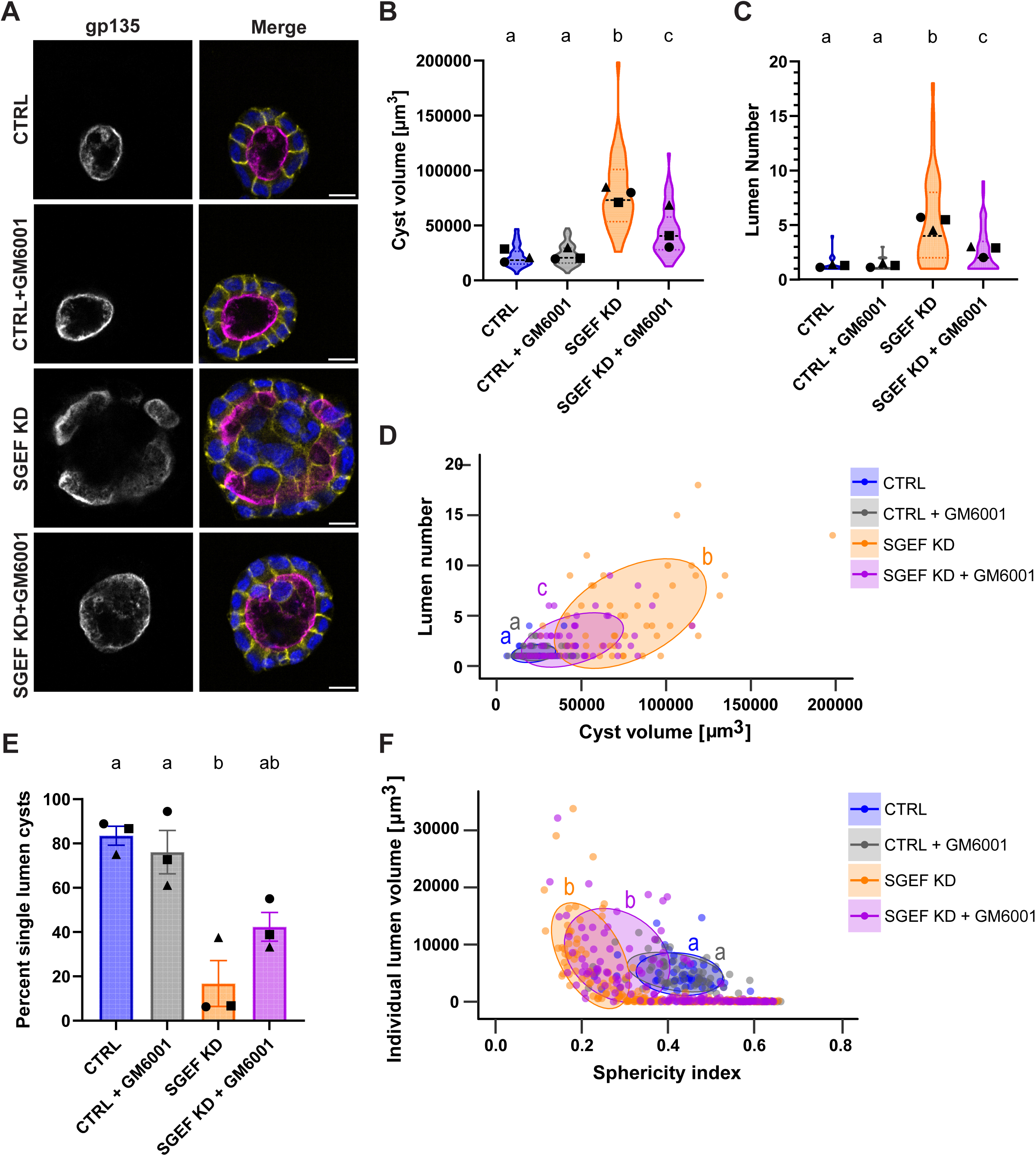
Metalloproteinase activity influences cyst volume and lumen phenotype in SGEF KD cysts. **(A)** Immunofluorescence of CTRL and SGEF KD cysts grown in the presence or absence of GM6001 and stained for gp135 (magenta), β-catenin (yellow), and Hoechst (blue). Scale bars: 10 µm. **(B)** Cyst volume. **(C)** Lumen number per cyst. **(D)** Relationship between lumen number and cyst volume. Each point represents one cyst. Ellipses include 60% of data in each condition. PERMANOVA was used for statistically comparing populations between ellipses. Letters indicate statistical groupings. **(E)** Percentage of cysts with a single lumen. **(F)** Individual lumen volume versus sphericity. Each point represents one lumen. Ellipses include 60% of lumens > 1000 µm^3^ per condition. All cysts associated with this figure were cultured for 7 days. For (B-E), n=3 independent experiments (>15 cysts/condition/experiment). For (F), n=3 (>15 lumens/condition/experiment). Violin plots show data distribution; symbols represent experiment means. Center line, median; bounds, IQR. Conditions not sharing a letter are significantly different (P<0.05, Dunn’s test or PERMANOVA).

Despite rescuing lumen number, GM6001 did not restore normal lumen morphology. Lumen sphericity was highly variable and was not significantly different from untreated SGEF KD cysts (Fig. 8F). Analysis of individual lumens suggested that many remained large and partially collapsed, although a subset exhibited a more spherical morphology. Together, these findings suggest that MMP activity promotes cyst motility, fusion, and multi-lumen formation following SGEF depletion, whereas restoration of normal lumen architecture likely requires proper junctional organization.

## Discussion

The establishment and maintenance of a polarized epithelium require the coordinated regulation of cell polarity, cell-cell adhesion, cytoskeletal organization, and interactions with the extracellular matrix (Roignot et al., 2013). A key player in the establishment of epithelial polarity is the Scribble polarity complex, composed of Scribble, Dlg1, and Lgl, which functions as a scaffold to coordinate diverse signaling pathways involved in cell polarity, junction assembly, and epithelial barrier function (Stephens et al., 2018)(Elsum et al., 2012). However, even though the role of the Scribble complex in epithelial organization has been well established, the downstream signaling pathways that coordinate these processes during tissue morphogenesis remain incompletely understood.

Our previous work established that SGEF forms a ternary complex with Scribble and Dlg1 to regulate adherens and tight junction integrity in epithelial monolayers (Awadia et al., 2019; Rabino et al., 2024). In this study, we identify SGEF as a critical regulator of epithelial morphogenesis in three-dimensional epithelial cysts. By combining quantitative morphometric analyses with long-duration live-cell imaging, we demonstrate that loss of SGEF disrupts lumen architecture, weakens cell-cell junctions, alters actomyosin organization, promotes collective migration and cyst fusion, and ultimately results in the formation of enlarged, multi-lumen cysts. Together, our findings support a model in which SGEF integrates junctional stability, cytoskeletal organization, and extracellular matrix remodeling to preserve epithelial architecture during lumenogenesis.

Importantly, SGEF-deficient cysts retain overall apicobasal polarity, but they fail to generate or maintain a single spherical lumen. Instead, the luminal compartment becomes highly heterogeneous, consisting of multiple lumens that vary considerably in size and morphology. These observations suggest that SGEF is required to preserve the mechanical integrity of the epithelial sheet as luminal pressure increases during cyst development.

Several observations support the idea that altered epithelial architecture contributes to this phenotype. SGEF depletion reduced the expression of both E-cadherin and ZO-1, two proteins that are essential for maintaining epithelial cohesion and barrier function. In addition, junctions became more linear, and non-muscle myosin IIB was redistributed throughout the epithelium, a phenotype that has been associated with altered force transmission across the epithelial sheet (Choi et al., 2016; Tokuda et al., 2014). Together, these changes are expected to compromise the ability of neighboring cells to coordinate mechanical forces required for lumen expansion. Restoration of E-cadherin and ZO-1 expression through inhibition of β-catenin signaling substantially improved lumen morphology and junction organization, demonstrating that disruption of epithelial junctions is a major contributor to the morphogenetic defects observed following SGEF depletion. However, β-catenin inhibition only partially rescued the multi-lumen phenotype, indicating that impaired junctional stability alone cannot fully explain the consequences of SGEF loss.

One of the most unexpected findings associated with SGEFKD was the remarkable increase in cyst motility. While CTRL MDCK cysts displayed limited displacement throughout development, SGEF KD cysts remained highly motile, frequently extending groups of cells while maintaining cell-cell contacts and exhibiting characteristics of collective migration rather than complete EMT. Restoring E-cadherin expression was sufficient to completely suppress this migratory behavior, indicating that reduced epithelial cohesion is a key driver of cyst motility. These observations further support previous work identifying E-cadherin loss as a central determinant of epithelial cell migration and suggest that SGEF normally functions to stabilize epithelial architecture by maintaining adhesive interactions between neighboring cells (Theveneau & Mayor, 2012). Our results also showed that SGEF KD cysts were significantly larger in volume. We initially thought that it was due to increased proliferation, but live-cell imaging revealed that the increased migration observed in SGEF KD cysts was closely associated with an increased frequency of fusion between neighboring cysts. This increase in motility and fusion, together with the observation that inhibition of matrix metalloproteinases restored cyst volume to CTRL levels, suggests that increased cyst volume in the absence of SGEF results primarily from fusion rather than enhanced proliferation. We also explored the possibility that multi-lumen cysts were a result of fusion of developing cysts. However, our results suggest that SGEF KD cysts develop multiple lumens even in the absence of fusion, and fusion just exacerbates this phenotype.

The observation that inhibition of matrix metalloproteinases substantially rescued both cyst volume and lumen number was particularly intriguing. Matrix metalloproteinases are well-established regulators of collective migration, branching morphogenesis, and tubulogenesis, where they remodel the surrounding extracellular matrix to facilitate tissue reorganization (Page-McCaw et al., 2007). Their contribution to epithelial lumen morphogenesis has received comparatively less attention, but there are some examples of MMPs role in lumen morphogenesis, mammary epithelial branching, and endothelial tubule formation (Simian et al., 2001; Singh et al., 2021; Stratman et al., 2009; Weaver et al., 2014). Surprisingly, inhibition of metalloproteinases rescued lumen number more effectively than inhibition of β-catenin signaling, suggesting that extracellular matrix remodeling represents a parallel pathway contributing to the SGEF phenotype rather than simply acting downstream of junction destabilization.

Our previous studies demonstrated that the catalytic activity of SGEF is required for maintaining E-cadherin and ZO-1 expression and for normal lumen formation (Awadia et al., 2019; Rabino et al., 2024), suggesting that these phenotypes are mediated through activation of RhoG. However, restoring catalytically active SGEF to the basolateral membrane in the absence of the Scribble/SGEF/Dlg1 ternary complex fails to rescue all the defects associated with SGEF depletion (Rabino et al., 2024). These findings suggest that SGEF contributes to epithelial morphogenesis through both its catalytic activity and scaffolding functions that depend on its association with the Scribble polarity complex.

One attractive possibility is that SGEF regulates epithelial architecture by coordinating multiple pathways centered on the stability of epithelial junctions. E-cadherin and ZO-1 are not only structural components of adherens and tight junctions but also function as mechanosensory platforms that couple cell-cell adhesion to the actomyosin cytoskeleton (Lecuit & Yap, 2015; Priya & Yap, 2015; Zihni et al., 2016). Loss of either protein has previously been associated with multi-lumen cyst formation and altered epithelial morphogenesis (Desclozeaux et al., 2008; Jia et al., 2011; Odenwald et al., 2017), while ZO-1 has recently been shown to regulate lumen architecture by coordinating junctional tension with hydrostatic pressure within the lumen (Mukenhirn et al., 2024). We previously demonstrated that SGEF depletion destabilizes the E-cadherin-catenin complex, promoting E-cadherin internalization and β-catenin release from adherens junctions. β-catenin signaling subsequently induces expression of the transcriptional repressor Slug, leading to further repression of E-cadherin transcription (Rabino et al., 2024). Together with the findings described here, these observations support a model in which disruption of epithelial junctions represents one of the earliest events following loss of SGEF, initiating a cascade of changes in tissue mechanics that ultimately impair lumen morphogenesis. Loss of epithelial cohesion is clearly an important determinant of cyst migration; however, additional signaling pathways are also likely to contribute. The Scribble complex has long been implicated in establishing front-rear polarity during directed migration, while both RhoG and its downstream signaling pathways regulate cell adhesion, lamellipodia formation, and cell migration (Abedrabbo & Ravid, 2020; Hiramoto-Yamaki et al., 2010; Hiramoto et al., 2006; Katoh et al., 2006; Meller et al., 2008; Osmani et al., 2006; Zinn et al., 2019). In particular, activated RhoG interacts with its downstream effector ELMO (engulfment and cell motility), which forms a complex with the Rac-GEF DOCK1 to promote localized Rac1 activation (Brugnera et al., 2002; Katoh & Negishi, 2003). Interestingly, ELMO2 has also been implicated in the recruitment and recycling of E-cadherin to nascent cell-cell junctions through Rab11-dependent trafficking and its interaction with integrin-linked kinase (Ho et al., 2016; Toret et al., 2014; Toret et al., 2018). These observations raise the intriguing possibility that the SGEF-RhoG signaling axis coordinates both junction assembly and Rac-dependent cytoskeletal remodeling through ELMO2, thereby integrating epithelial adhesion with collective cell migration.

The mechanism by which SGEF regulates metalloproteinase activity remains unknown. β-catenin signaling has been shown to induce MT1-MMP expression in invasive cancer cells, whereas intact cadherin/catenin complexes in epithelial and endothelial cells, keep β-catenin signaling low, which results in low MT1-MMP expression (Doyle & Haas, 2009; Kiran et al., 2011; Liu et al., 2010). However, MMP expression and activation can also be transcriptionally regulated independently of β-catenin (Yan & Boyd, 2007), and post-transcriptionally in response to cytoskeletal remodeling and Rho GTPase signaling (Ispanovic & Haas, 2006; Ispanovic et al., 2008). Consistent with this possibility, we previously demonstrated that depletion of either SGEF or RhoG enhances ECM degradation and invasion (Goicoechea et al., 2017). Together with our current findings, these observations suggest that SGEF normally suppresses extracellular matrix remodeling in both normal epithelial tissues and invasive tumor cells. Whether this regulation occurs through transcriptional control of metalloproteinases, altered trafficking of membrane-associated proteases such as MT1-MMP, or indirect effects on cytoskeletal organization remains an important question for future investigation.

Collectively, our results support a model in which SGEF functions as a signaling hub that coordinates epithelial adhesion, cytoskeletal organization, and extracellular matrix remodeling during epithelial morphogenesis. We propose that loss of SGEF initially destabilizes adherens and tight junctions, reducing epithelial cohesion and altering actomyosin organization. These changes not only impair the mechanical integrity required for lumen expansion but also promote collective migration and extracellular matrix remodeling, leading to cyst fusion and persistence of abnormal lumen architecture. Rather than acting exclusively through RhoG activation, SGEF appears to function within the Scribble polarity complex to spatially integrate multiple signaling pathways required for epithelial tissue organization. Elucidating how SGEF coordinates these diverse processes should provide important insight into the mechanisms by which epithelial polarity is maintained during normal development and disrupted during pathological conditions such as carcinoma progression.

## Materials and Methods

### Cell lines

MDCK II T23 cells were a gift from Ian Macara (Vanderbilt University, Nashville, TN). They were grown in high-glucose DMEM supplemented with L-glutamine and sodium pyruvate (Cytiva), 10% fetal bovine serum (FBS, Cytiva), and penicillin–streptomycin (Corning). Cells were incubated in a humid environment at 37°C with 5% CO_2_. All experiments were conducted with cells that were passaged under 30 times. Mycoplasma contamination was routinely tested by PCR amplification. CTRL, SGEF KD, WT Rescue, and SGEF/Slug KD MDCK stable cell lines were described previously (Awadia et al., 2019; Rabino et al., 2024).

### MDCK 3D cysts

The MDCK cyst culture protocol was adapted from Anirban Datta and David Bryant (Mostov Lab, UCSF, San Francisco, CA). One day prior to cyst plating, maintenance cultures were subcultured at a 1:8 ratio to reduce confluency and allow preparation of a single-cell suspension.

Growth Factor Reduced Matrigel aliquots (354230, Corning) were thawed on ice and used to coat µ-Slide 8 well glass bottom chamber slides (80807, Ibidi) with 15 µL of 100% Matrigel per well, followed by polymerization at 37°C for approximately 30 minutes.

MDCK cells were detached using 0.25% trypsin, neutralized with complete medium (high glucose DMEM supplemented with 10% FBS and 1% penicillin-streptomycin), counted, and resuspended at 3,000 cells per well. Cells were mixed at 1:1 with a 4% Matrigel solution prepared in complete medium to achieve a final concentration of 2% Matrigel and plated in a final volume of 300 µL per well. Medium containing 2% Matrigel was replaced every other day. Cysts were cultured for 5-7 days to allow polarization and lumen formation before fixation for immunofluorescence.

### Antibodies and reagents

For IF of MDCK cysts, primary and secondary antibodies were used at twice the concentration typically applied for 2D monolayers. All IF dilutions listed describe use for 3D cysts. For Western blot analysis of cyst samples, antibodies were used at the same concentrations as for 2D Western blots. However, membranes were incubated with primary antibodies for at least 1.5 hours at room temperature. For some proteins such as ZO-1, primary antibody incubation was performed overnight at 4°C.

The following commercial antibodies were used in this study: anti-tubulin (T9028, mouse monoclonal; Sigma-Aldrich) 1:40,000 (WB); anti-E-cadherin (24E10, rabbit mAb; Cell Signaling Technology) 1:1,000 (WB), 1:75 (IF); anti-Slug (C19G7, rabbit mAb; Cell Signaling Technology) 1:500 (WB); anti-myosin IIB (D8H8 XP, rabbit polyclonal; Cell Signaling Technology) 1:100 (IF); anti-phospho-myosin light chain 2 (Ser19, 3675S, mouse mAb; Cell Signaling Technology) 1:75 (IF); anti-gp135 (3F2/D8, mouse mAb; DSHB) 1:200 (IF); anti–ZO-1 (R26.4C-s, rat IgG1; DSHB) 1:15 (IF); anti–mNeonGreen (32f6, mouse mAb; Proteintech) 1:250 (IF); anti-mNeonGreen (CTK0203, rabbit mAb; Chromotek) 1:500 (IF); anti-β-catenin (A302-010A, rabbit IgG; Bethyl Laboratories) 1:250 (IF), 1:1,000 (WB); anti–ZO-1 (339100, mouse mAb; Thermo Fisher Scientific) 1:1,000 (WB), 1:100 (IF). Secondary antibodies for IF included Alexa Fluor 488, 568, and 647-conjugated anti-mouse, anti-rabbit, and anti-rat IgG antibodies (A11008, A11001, A11005, A32733, R37117; Thermo Fisher Scientific) used at 1:500, and Alexa Fluor 647-conjugated phalloidin (A22287; Thermo Fisher Scientific) used at 1:750. HRP-conjugated anti-mouse and anti-rabbit secondary antibodies (715-035-151, 711-035-152; Jackson ImmunoResearch) were used at 1:20,000 for Western blotting. Hoechst 34580 (63493-51G; Sigma) was used 1:750 in IF to stain nuclei.

The following reagents were also used throughout this study. For β-catenin inhibition, iCRT3 (219332; EMD Millipore) was used at 50 µM. For MMP inhibition, GM6001 (14533; Cayman Chemical) was used at 25 µM.

### Constructs

The LV-GFP H2B construct (Addgene 25999) was a gift from Dr. Joachim Goedhart (Swammerdam Institute for Life Sciences, Amsterdam, The Netherlands). The pLenti-CAAX Scarlet construct was a gift from Dr. Jaap van Buul (University of Amsterdam, Amsterdam, The Netherlands).

### Lentiviral particles and infection

Lentiviral particles were packed in HEK293FT cells using a standard calcium phosphate transfection protocol as described previously in (Awadia et al., 2019). Briefly, HEK293FT cells were transfected with 2.2 µg of pMD2.G, 4 µg of pSPAX2, and 3.735 µg of the lentivirus plasmid. The pMD2.G and pSPAX2 packaging plasmids were a gift from Didier Trono, EPFL, Lausanne, Switzerland (Addgene plasmids 12259 and 12260). The culture medium was changed 24 h after transfection, and lentivirus particles were harvested 48 h after transfection.

MDCK cells were infected overnight with lentivirus particles along with polybrene (1:1000). The following day, the infection medium was replaced with complete growth medium. For some viruses, single-cell colonies were isolated by serial dilution and examined under confocal microscopy for the desired fluorescence. Scarlet CAAX/GFP H2B double-expressing cells were enriched by fluorescence-activated cell sorting (FACS) at the University of Toledo Flow Cytometry Core Facility (Toledo, OH).

### SDS-PAGE and Western blotting of 3D cysts

Cysts cultured for 7 days at a starting density of 6.25×10^4^ per well in a 12-well dish were rinsed with ice cold PBS three times and then scraped into 100 µL lysis buffer containing 150 mM NaCl, 0.5 mM MgCl_2_, 0.2 mM EGTA, 50 mM Tris-HCl pH 7.4, 1% triton-X100, and EZBlock protease inhibitor cocktail (BioVision). Samples were then transferred to a microcentrifuge tube, incubated on ice for 15 minutes and sonicated for 10 seconds at 30% power. Lysates were then centrifuged at 4°C at 13,000 rpm for 15 min. The supernatant was then transferred to a new, pre-chilled microcentrifuge tube. For Western blotting, lysates were boiled in 2X Laemmli buffer, and 40-60 μg of protein were resolved by SDS-PAGE. The proteins were transferred onto PVDF membranes and immunoblotted with the indicated antibodies.

Immunocomplexes were visualized using the SuperSignal West Pico PLUS Chemiluminescent HRP substrate (Thermo Fisher Scientific).

### Immunofluorescence assays and microscopy

The IF assay protocol was adapted from Anirban Datta and David Bryant from the Mostov Lab (UCSF, San Francisco, CA). MDCK cysts grown for 5-7 days in µ-Slide 8 well glass bottom chamber slides (Ibidi) were fixed for 15 min with 4% paraformaldehyde and quenched for 10 minutes with 10 mM ammonium chloride. Cysts were then washed three times with PBS, and permeabilized with 0.1% Triton X-100 in PBS for 10 min. The cysts were then washed with PBS three times, and blocked with a solution of PBS, 2.5% goat serum, and 0.2% Tween 20 for 20 minutes, followed by 20 minutes of blocking with a solution of PBS, 0.4% fish skin gelatin, and 0.2% Tween 20. Cysts were then incubated overnight at 4°C with the primary antibody. The following morning, cysts were washed three times for 5 minutes and three times for 10 minutes with a solution of PBS and 0.2% Tween 20, followed by 20 minutes of blocking with PBS, 0.4% fish skin gelatin, 0.2% Tween 20, and 20 min with PBS, 2.5% goat serum, 0.2% Tween 20. The secondary antibodies, along with Hoechst, were diluted in the goat serum blocking solution and added to the cysts for 2-3 hours at room temperature or overnight at 4°C. They were then washed three times for five minutes and three times for ten minutes with PBS, 0.2% Tween, and mounted with a 1:3 mixture of CitiFluor AF200 (17977-10, Electron Microscopy Sciences) and CitiFluor MWL4-88 (17977-150, Electron Microscopy Sciences) respectively.

Most confocal images were acquired using an Andor Dragonfly 200 spinning disk confocal microscope (Oxford Instruments) mounted on a Leica DMi8 microscope stand and equipped with 20x/0.40 NA dry, 40x/0.95 NA dry, and 63x/1.40 NA oil immersion objectives, 4 laser lines (405, 488, 561 and 637 nm), and an Andor Zyla SCMOS camera. For live imaging, we used an Ibidi stage top incubator (set to 37°C and 5% CO2). For selected experiments we used an Andor Benchtop spinning disk confocal microscope equipped with a 60x/1.42 NA oil immersion objective and 4 laser lines (405, 488, 561 and 637 nm).

### Image analysis

All image analysis scripts were developed to process Imaris files but can be adapted for other file formats. The codes, as well as detailed instructions for each quantification workflow, are available on our GitHub repository (https://github.com/Garcia-Mata-Lab/cyst_quantification/tree/main. All workflows use a combination of Fiji/ImageJ and RStudio. For experiments with multiple quantifiable channels in the same cyst, the provided ImageJ macros and RStudio codes can be combined as needed. All ImageJ macros were used in batch processing format.

Cyst volume was quantified using the nuclear channel. In Fiji/ImageJ, a gaussian blur filter was applied, followed by a heavy threshold to define the cyst boundary. The “Fill Holes” function was used to eliminate internal gaps within the thresholded region. The “3D Objects counter” tool was then used to extract volume and surface area measurements for the cyst. In RStudio, volume measurements were filtered to remove measurements taken of background particles, and the final cyst volumes were exported to a CSV file.

Lumen morphology quantification was performed using the gp135 channel following a similar Fiji/ImageJ workflow as described for cyst volume, except with a more stringent threshold to preserve individual lumen architecture. In RStudio, objects less than 5 μm^3^ in volume were filtered from further analysis. The remaining lumen measurements were then tallied per cyst to obtain lumen number. The lumen sphericity index was calculated with the formula: *[³√(36·π· Vℓ ²) / s]* where Vℓ is lumen volume and *s* is lumen surface area (Wadell, 1932).

Quantification of cell number was obtained using a workflow integrating Fiji/ImageJ, Napari, and RStudio. Imaris files were converted in Fiji/ImageJ to single-channel TIFF stacks containing only the nuclear signal. Nuclei were then segmented throughout the Z-stack in Napari using the *napari-zelda* plugin, incorporating multiple thresholding parameters. In RStudio, segmented objects were filtered by volume to remove debris and apoptotic cells included during Napari segmentation. The remaining objects were then tallied to obtain the number of nuclei per cyst.

ZO-1, E-cadherin, and β-catenin intensities were quantified using a custom background subtraction macro in Fiji/ImageJ (Kreider-Letterman et al., 2023). The channel of interest was first smoothed using a Gaussian blur, and a predetermined threshold was applied to generate a binary mask of areas in the cyst containing signal. The binary mask was then multiplied by the original image, preserving the intensity values. A second predetermined threshold was applied to exclude any residual signal or autofluorescence. This threshold was adjusted for each protein and experiment and was not altered between conditions. All pixels below the threshold were set to NaN to eliminate background from being measured. The integrated density was measured in pixel units for each Z-slice, and in RStudio, these values were added across all slices for each cyst. To normalize the data to cell number, the total integrated density of each cyst was divided by the number of nuclei within that cyst [(Σ IntDen) / N].

Quantification of total non-muscle myosin IIB and p-MLC intensity and localization was done by first creating a sum projection of the 5 center slices of the cyst using the myosin or p-MLC channel in Fiji/ImageJ. A binary mask of the entire cyst was generated, and then the *Analyze Particles* function was used to extract the centroid and Feret’s diameter of the cyst. A point ROI was placed at the centroid, and a horizontal line ROI corresponding to the Feret’s diameter was drawn to span the longest axis of the cyst. By adapting a radial analysis macro previously created by our lab, this line ROI was rotated 360° at 1⁰ intervals. The resulting intensity measurements were averaged across all angles to generate a one-dimensional radial intensity profile extending from the cyst center toward the periphery (Kreider-Letterman et al., 2023). In R, to account for the line extending farther than the cyst boundary, each profile was automatically truncated based on a defined intensity threshold and interpolated to a standardized radial distance. Profiles were normalized to the peak fluorescence intensity within each cyst and subsequently scaled relative to the average peak intensity of the CTRL condition to preserve differences in absolute protein abundance between experimental groups. Cysts exhibiting extreme deviations from the mean profile of their respective condition were identified by Z-score analysis and excluded as outliers. Mean normalized intensity profiles +/- SEM were then calculated for each condition across biological replicates.

To quantify the zig-zag index, a max intensity projection of the ZO-1 channel covering only the most basal layer of the tight junction network was generated in Fiji/ImageJ. Two diagonal lines were drawn into an “X” spanning the entire image, and junctions contacting the X were measured to avoid bias. Compressed junctions along the edge of the lumen were not included in measurements. Each junction was traced manually using the freehand line tool to capture the actual curvature. A straight line was then drawn between the same start and end points using the straight-line tool to obtain the linear distance. The zig-zag index for each junction was calculated in Excel as the ratio of the traced length divided by the straight-line length.

To track cyst motility, max intensity projections of all z-slices were generated for each FOV in Fiji/ImageJ. A Gaussian blur with a radius of 2 was applied to reduce noise prior to tracking. Cysts were then tracked using the TrackMate plugin with a thresholding detector, which varied between FOVs. The LAP tracker was then used with these settings: frame to frame linking max distance 40 μm, segment gap closing max distance 35 μm with a max frame gap of 5, segment splitting max distance 15 μm, and segment merging max distance 15 μm.

This pipeline was used to obtain quantitative measurements of the migratory characteristics for multiple cysts concurrently in each FOV.

### Statistical Analysis

In this study, technical replicates were defined as individual cysts, lumens, or 2D cells.

Biological replicates were defined as the mean values of technical replicates obtained from independent experiments. Western blot experiments included only biological replicates. Error bars represent the standard error of the mean (SEM) unless otherwise specified.

All statistical analyses were done in GraphPad Prism and RStudio. Data was tested for normality using the Shapiro-Wilk test. For abnormally distributed data, either Welch’s t-test (two conditions) or a Kruskal-Wallis test with Dunn’s post-hoc multiple comparisons (more than two conditions) were applied. For normally distributed data, a Student’s t-test (two conditions) or Brown-Forsythe and Welch ANOVA with Dunnett’s T3 multiple comparisons test (more than two conditions) was used. Statistical significance was defined as P < 0.05 (*), P < 0.01 (**), P < 0.001 (***), and P < 0.0001 (****).

For multidimensional scatter plot analyses, differences in overall profile distributions between experimental groups were assessed using PERMANOVA in R. group separation was visualized using 60% confidence ellipses. Statistical differences between the distributions represented by the confidence ellipses were assessed using a permutation-based test, and P values were adjusted for multiple comparisons using the Benjamini-Hochberg false discovery rate (FDR) correction.

## Acknowledgements

The authors would like to thank Drs. Ian Macara, Joachim Goedhart, and Jaap van Buul for providing reagents, Dr. Wei Niu for her support on using the Andor benchtop confocal, and all members of the Garcia-Mata lab for helpful discussions. This work was supported by grants from the National Institutes of Health to R.G.-M. (R01GM136826 and R15GM155874). FACS analysis was supported by NIH grant 1S10OD036266-01 for the BD FACS Discover S8 Cell Sorter, housed in the University of Toledo Flow Cytometry Core Facility (University of Toledo, Toledo, OH).

## Supplemental Figure Legends

**Supplementary Figure 1.**
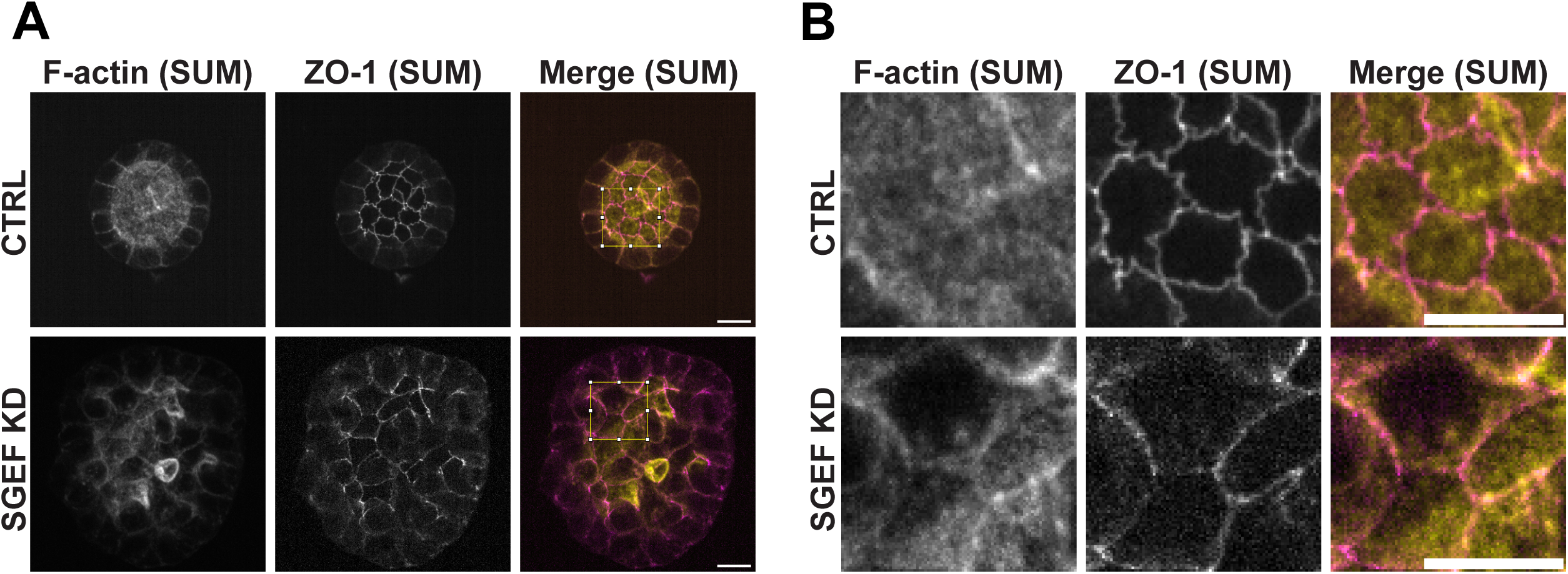
F-actin and ZO-1 localization in MDCK cysts. Representative immunofluorescence of day 7 CTRL and SGEF KD cysts stained for F-actin (yellow) and ZO-1 (magenta). **(B)** Zoomed images of boxed areas in (A). Images are sum projections (SUM) of the basal-most z-slices containing the tight junction network. Scale bars: 10 µm.

**Supplementary Figure 2.**
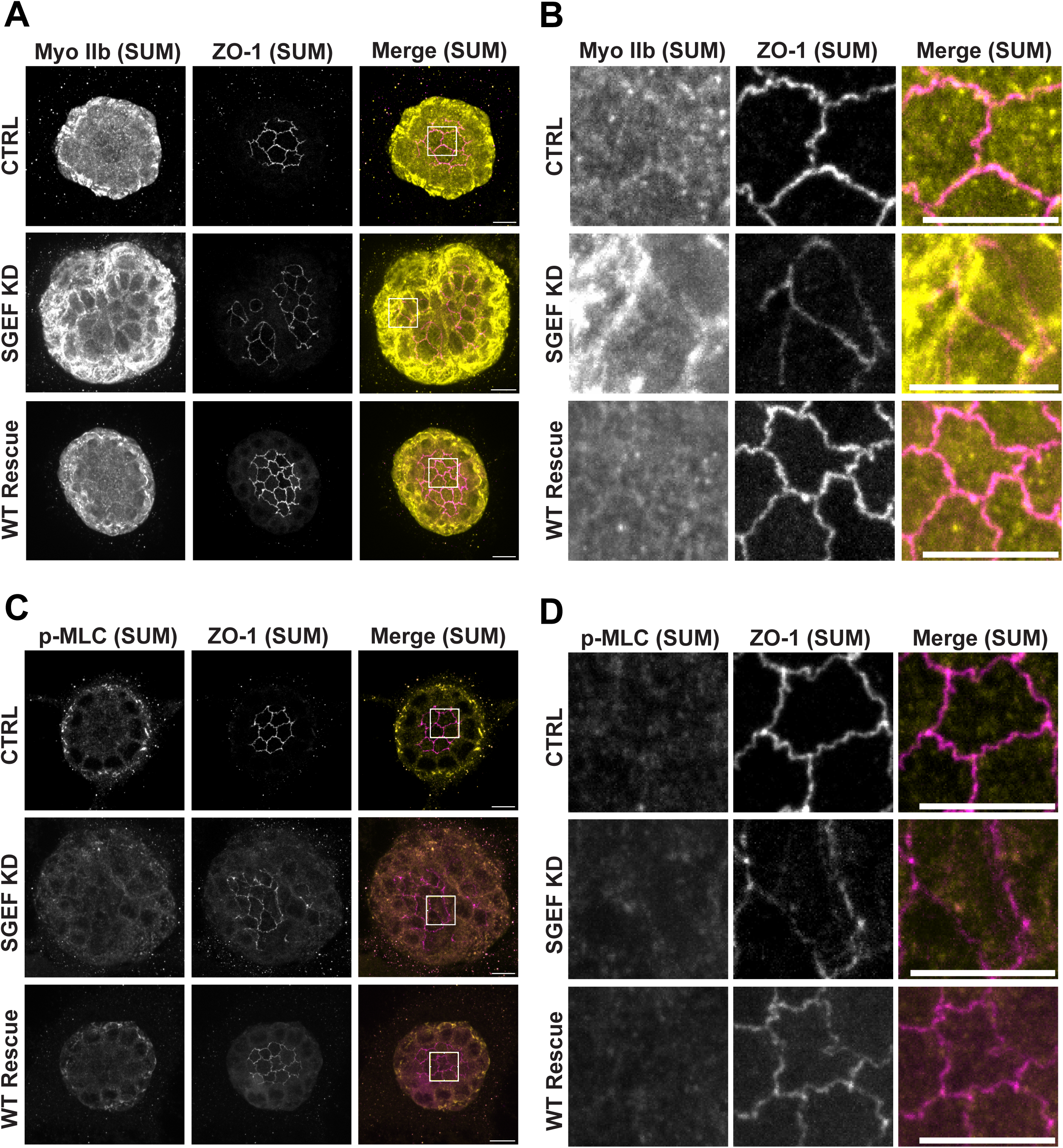
Myosin localization at tight junctions. **(A)** Representative immunofluorescence of day 7 CTRL, SGEF KD, and WT Rescue cysts stained for non-muscle myosin IIB (yellow) and ZO-1 (magenta). **(B)** Zoomed images of boxed areas in (A). **(C)** Representative immunofluorescence stained for p-MLC (yellow) and ZO-1 (magenta). **(D)** Zoomed images of the boxed areas shown in (C). All images are sum projections (SUM) of the basal-most z-slices containing the tight junction network. Scale bars: 10 µm.

**Supplementary Figure 3.**
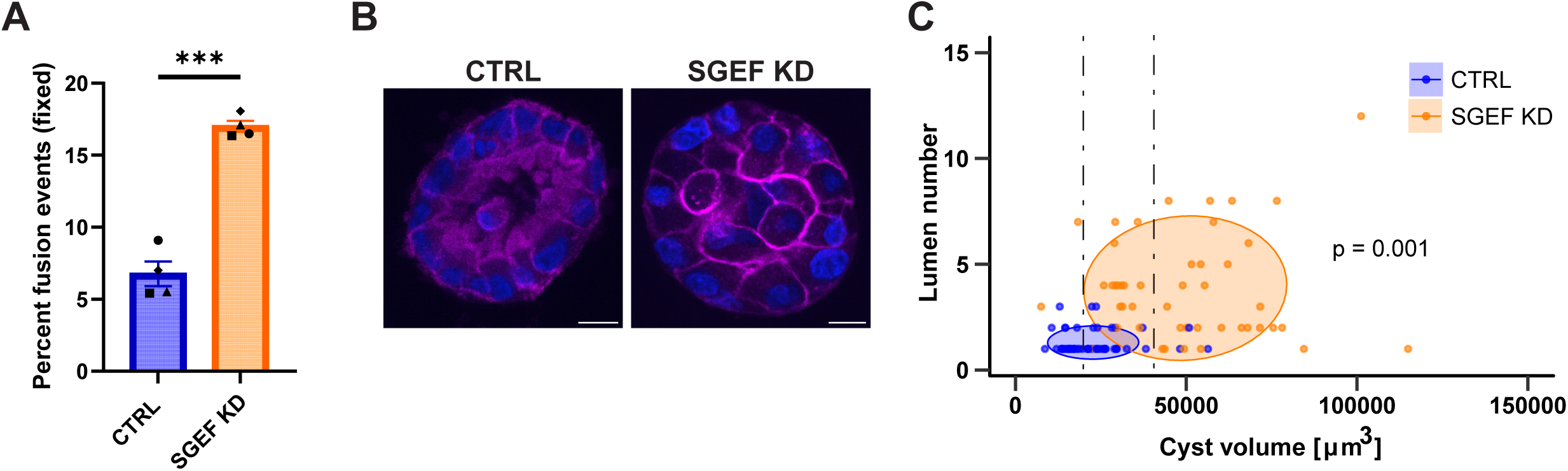
Cyst fusion is not required for the multi-lumen phenotype. **(A)** Percentage of total cysts from (Fig. 6A) that were dually labeled with Scarlet-CAAX and GFP-H2B, to detect fusion events between at least two cysts. Symbols represent experiment means (n=4 independent experiments; ∼80 cysts/condition/experiment). *\*\*\*P*<0.001. **(B)** Representative images of non-fused, similarly sized CTRL and SGEF KD Scarlet CAAX (magenta) cysts fixed after 84 hours of live imaging. Cysts were stained with Hoechst (blue) to visualize nuclei after fixation. Scale bars: 10 µm**. (C)** Relationship between lumen number and cyst volume for individual cysts. Ellipses encompass 60% of the data (n=3; >15 cysts/condition/experiment). PERMANOVA results are represented by the p-value displayed on the graph.

**Video 1. Three-dimensional reconstruction of lumen architecture. (A)** Imaris segmentation and 3D reconstruction of the single, central lumen in a representative day 5 CTRL cyst labeled with gp135. **(B)** Reconstruction of multiple lumens in a representative SGEF KD cyst. Videos show a 360° rotation along the vertical axis.

**Video 2. Three-dimensional reconstruction of tight junction network architecture.** Imaris segmentation and 3D reconstruction of ZO-1-labeled tight junction networks in representative day 7 **(A)** CTRL, **(B)** SGEF KD, and **(C)** WT Rescue cysts. Videos show 180° boomerang rotations along the vertical axis.

**Video 3. iCRT3 partially restores tight junction architecture.** Imaris segmentation and 3D reconstruction of ZO-1-labeled tight junction networks in representative day 7 **(A)** SGEF KD and **(B)** SGEF KD + iCRT3 cysts. Videos show 180° boomerang rotations along the vertical axis.

**Video 4. Live imaging of cyst migration and fusion.** 20x objective live imaging of Scarlet-CAAX-expressing CTRL and SGEF KD cysts from 12-59.5 hours after seeding, showcasing the period of greatest migration. 50-µm z-stacks were acquired every 30 minutes and displayed as max projections. Time after seeding is shown in the upper left. Scale bars: 40 µm. **(A)** Representative CTRL cysts exhibiting cell division and rotation without migration or fusion **(B)** Representative SGEF KD cysts showing asymmetric behavior, migration, and fusion.

**Video 5. E-cadherin expression suppresses SGEF KD cyst migration.** 20x objective live imaging of SGEF KD and SGEF/Slug KD cysts stably expressing Scarlet-CAAX from 12-59.5 hours after seeding. 50-um z-stacks were acquired every 30 minutes and displayed as max projections. Time after seeding is shown in the upper left. Scale bars: 40 µm. **(A)** Representative SGEF KD cysts showcasing extensive migration and fusion. **(B)** Representative SGEF/Slug dKD cyst showing marked suppression of migration, despite some persistence of abnormal cell rearrangements during rotation.

## Notes

### Competing Interest Statement

The authors have declared no competing interest.

